# Sexually dimorphic behavioural signatures of tau toxicity in adult *Drosophila*

**DOI:** 10.64898/2026.06.22.733697

**Authors:** Edmond N Mouofo, Maxwell P Spires-Jones, Yu-Chun Wang, Nils Schoovaerts, Patrik Verstreken, Claire S Durrant, James H Catterson, Tara L Spires-Jones

**Affiliations:** Institute for Neuroscience and Cardiovascular Research | The University of Edinburgh | Edinburgh | UK; UK Dementia Research Institute | The University of Edinburgh | Edinburgh | UK; VIB Center for Neuroscience Leuven, Belgium; Department of Neurosciences | Leuven Brain Institute | KU Leuven | Leuven | Belgium

**Keywords:** *Drosophila*, Neurodegeneration, Tauopathy, Alzheimer’s disease, Tau, Behaviour, Sex, Sexual dimorphism, Vacuoles, Sleep

## Abstract

Tau pathology is central to Alzheimer’s disease and related tauopathies, yet mechanisms driving neuronal dysfunction and degeneration downstream of pathological changes in tau remain poorly understood. *Drosophila melanogaster* models provide a genetically tractable system with an intact nervous system and short lifespan that allows investigation of mechanisms of many diseases. However, in *Drosophila*, developmental expression of human tau frequently causes lethality and developmental phenotypes, limiting the study of neurodegenerative disease processes. Further, sex is rarely considered in *Drosophila* studies of tau pathology despite clear sex differences being observed in many aspects of human tauopathies. Here, we used an inducible, pan-neuronal GeneSwitch system to express human tau isoforms exclusively in adulthood, enabling the dissection of tau toxicity independent of development. We combined longitudinal behavioural monitoring with lifespan and neurodegeneration analyses, and performed a targeted genetic screen to identify modifiers of tau-induced dysfunction. Adult-onset tau expression produced striking, sexually dimorphic effects on survival and behaviour. Neuronal expression of the human tau isoform with 4 microtubule binding repeats and neither alternatively spliced N-terminal exon (0N4R tau) caused pronounced neurodegeneration and reduced lifespan, which was exacerbated in flies expressing the phospho-mimetic 0N4R-Tau^E14^ variant. Tau expression produced sexually dimorphic effects on survival and behaviour, with females exhibiting a greater reduction in lifespan, while the induction-dependent increase in vacuolar neurodegeneration was broadly comparable between sexes. Behaviourally, tau expression induced elevated daytime inactivity in females, whereas males exhibited hyperactivity, revealing opposing functional outcomes between sexes. A targeted genetic screen further identified modifiers of tau-dependent behavioural impairment. *APOE2* expression in glia, syndecan overexpression in neurons, and increased global expression of the chaperone heat shock protein 90 all reduced 0N4R-Tau^E14^-induced behavioural changes. Seventeen candidate perturbations enhanced the Tau^E14^-induced behavioural phenotype, including manipulations of APOE3, CLU, INPP5D/INPP5K, BIN1/Amph, synaptogyrin, LRP1, NPC1, and Hsp90 pathways. Together, these findings establish an adult-onset *Drosophila* model of tauopathy that uncouples neurotoxicity from development, reveals sex as a major determinant of tau-induced behavioural outcomes in flies, and uncovers genetic modulators of tau-induced dysfunction. This work highlights the importance of incorporating sex as a biological variable and provides a platform for mechanistic and translational studies of tauopathy.

## Introduction

Abnormal tau accumulation defines a spectrum of neurodegenerative disorders collectively termed tauopathies^1^. In these conditions, tau aggregation is closely associated with neuronal dysfunction and progressive neurodegeneration, with clinical features reflecting the brain regions affected^2,3^. Variants in the *MAPT* gene encoding tau cause neurodegenerative tauopathies, and in animal models, expressing human tau with disease-associated variants induces neurodegeneration, making tau a central causal factor rather than a passive disease marker^4^. Understanding how tau pathology disrupts neuronal function and causes neurodegeneration remains a critical challenge in the field.

*Drosophila melanogaster* provides a powerful system for addressing this challenge. The fly nervous system shares conserved cellular machinery with mammals, including pathways governing cytoskeletal organisation, synaptic transmission, and axonal transport^5^. Neuronal expression of human tau in *Drosophila* recapitulates key features of tau-mediated toxicity, including reduced lifespan, locomotor dysfunction, vacuolar neurodegeneration, synaptic abnormalities, and molecular hallmarks of tau pathology^6–10^. Together, these properties make *Drosophila* particularly well suited for dissecting tau-driven neurodegeneration.

Behavioural phenotyping has emerged as a sensitive means of detecting tau-induced neuronal dysfunction in *Drosophila*. Automated locomotor and sleep analyses using the *Drosophila* Activity Monitor (DAM) provide continuous, unbiased measures of activity, circadian rhythms, and sleep architecture^11,12^. Tau expression in flies has been shown to disrupt locomotion, weaken circadian rhythms, and alter sleep patterns, mirroring early non-cognitive symptoms observed in human tauopathies^5,13,14^. Notably, behavioural impairments can be detected before extensive neuronal loss or overt histopathological degeneration, highlighting their utility as early functional readouts of tau toxicity^13,15,16^.

Despite the extensive use of *Drosophila* models to study tau-induced neuronal dysfunction, sex has rarely been included as an experimental variable, despite known sex differences in human tauopathies including in disease prevalence, pathology, and response to drugs across several diseases ^17–20^. Several *Drosophila* studies have expressed human tau pan-neuronally or within specific neuronal populations to investigate neurodegeneration, neuronal physiology and behavioural phenotypes, including locomotor activity and sleep and circadian rhythms^13,21^. These behavioural analyses have typically not assessed sex as a factor and have used exclusively male flies^21–23^. Indeed, most published DAM-based behavioural analyses are performed in males - likely due to practical considerations, as females lay eggs that can hatch within monitoring tubes, introducing larvae that disrupt infrared beam crossings and limit the duration and reliability of recordings^24^. Additional variability associated with female reproductive state, along with historical precedent in the field, may further contribute to this bias^25,26^. While some studies report behavioural and physiological phenotypes following neuronal tau expression, including circadian disruption, hyperactivity and reduced sleep^13^, or isoform-specific neuronal dysfunction and lifespan effects^21^, sex-specific responses are not examined. Even more broadly, large-scale and systems-level studies of tau toxicity in *Drosophila* similarly do not incorporate sex-stratified analyses^27–29^.

As a result, the extent to which sex modulates behavioural outcomes in *Drosophila* tauopathy models remains largely unresolved. Here, we address this gap by explicitly examining female flies alongside males in DAM-based behavioural analyses of neuronal tau expression.

An important limitation of many existing *Drosophila* tau models is that tau expression is initiated during embryonic development, making it difficult to distinguish developmental abnormalities from adult-onset neurodegeneration. Conditional expression systems such as GeneSwitch-UAS overcome this limitation by enabling temporally controlled induction of tau expression in adult neurons^30,31^. Adult-onset tau expression therefore provides a framework for modelling age-associated neurodegenerative processes and for identifying early phenotypes that precede irreversible neuronal loss. By combining adult-onset neuronal tau expression with behavioural monitoring using DAMs, it is possible to detect early, progressive functional consequences of tau toxicity and link these to subsequent neurodegenerative outcomes.

In this study, we utilise the GeneSwitch-UAS system to induce adult-onset pan-neuronal expression of human tau forms in *Drosophila*, with the aim of developing a robust model of tau-induced neurodegeneration allowing the study of potential modifiers of tau toxicity.

## Materials and methods

### Fly husbandry and stocks

Fly maintenance and experiments were conducted at 25°C under a 12 h:12 h light:dark (LD) cycle in temperature-controlled incubators (Panasonic MIR254-PE). Fly food contained the following ingredients: water, agar (BTP Drewitt, 960), brewer’s yeast (BTP Drewitt, AYP), glucose (Fisher Scientific, 11472858), maize (MP Biomedicals, 11455522), heat-killed Fermipan yeast (autoclaved at 116°C for 15 min and stored as stock) (Fermipan Red Yeast 500g (Lallemand, ASIN B07K91GYJG), Amazon), propionic acid (Thermo Fisher 149300025), and Nipagin (BTP Drewitt, Nipam-B1-O1NP) pre-dissolved in ethanol. Adult-onset, pan-neuronal transgene expression was induced using *Elav-GeneSwitch* (*Elav-GS*) by supplementing food with RU486 (mifepristone; Sigma-Aldrich, M8046-1G) to a final concentration of 200 µM. The full list of genotypes used in this study are shown in **Table 1**.

**Table 1.**
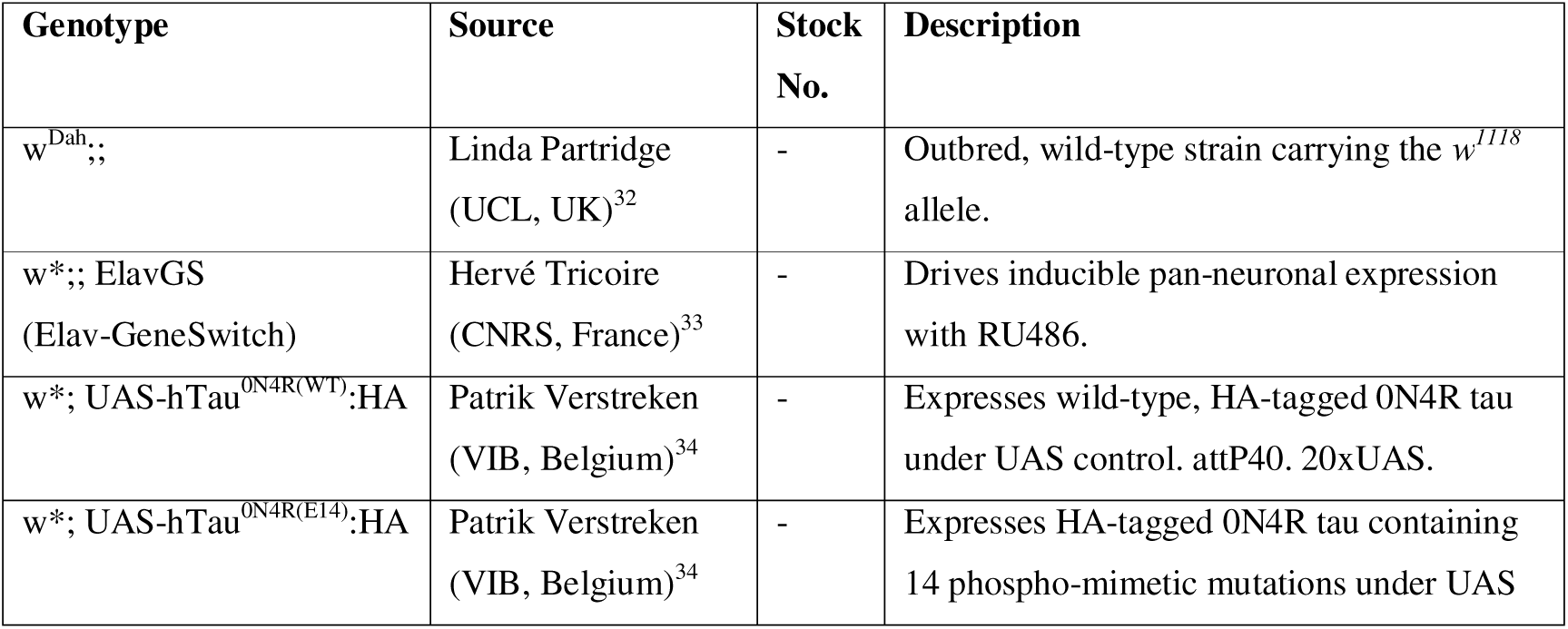
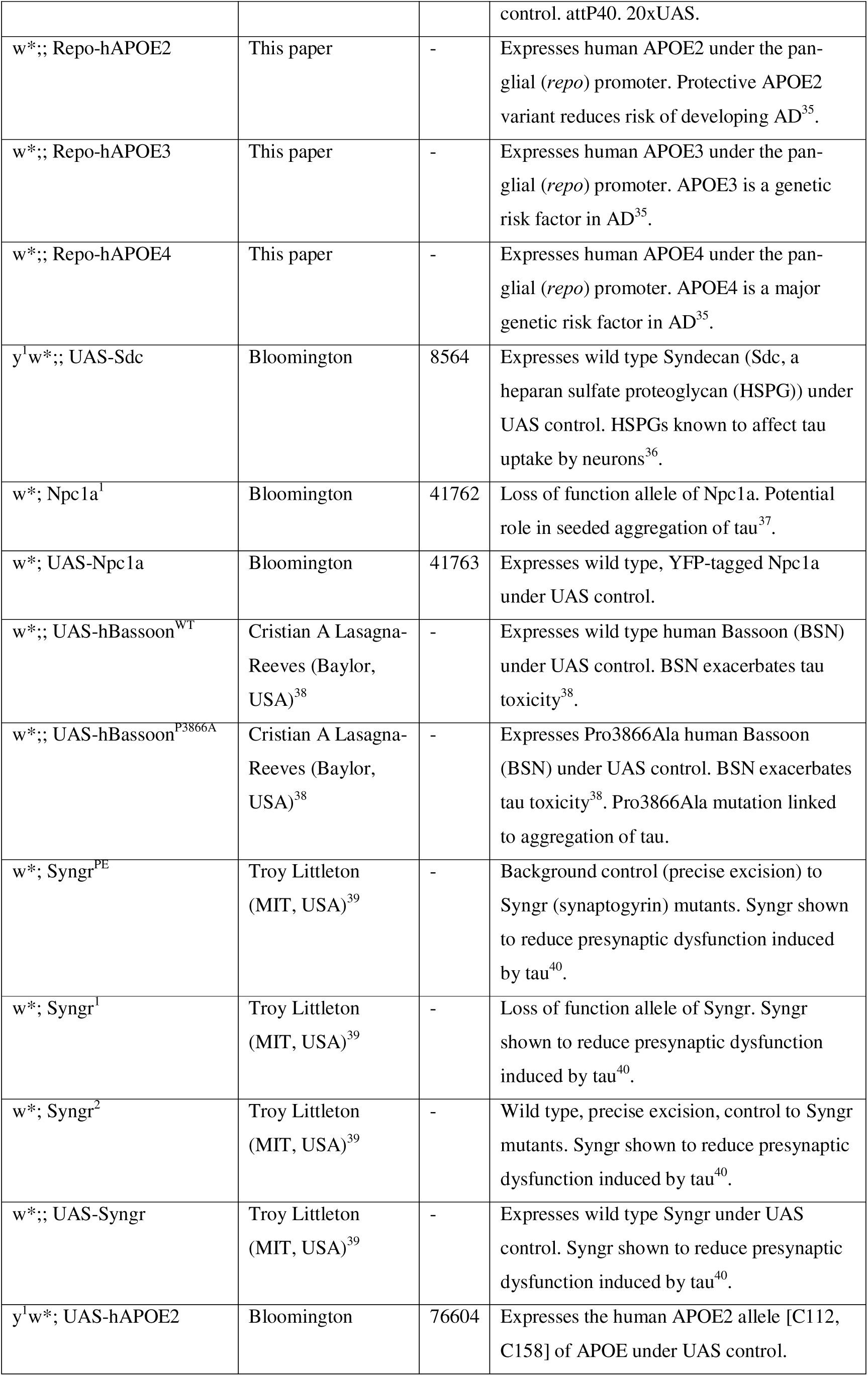
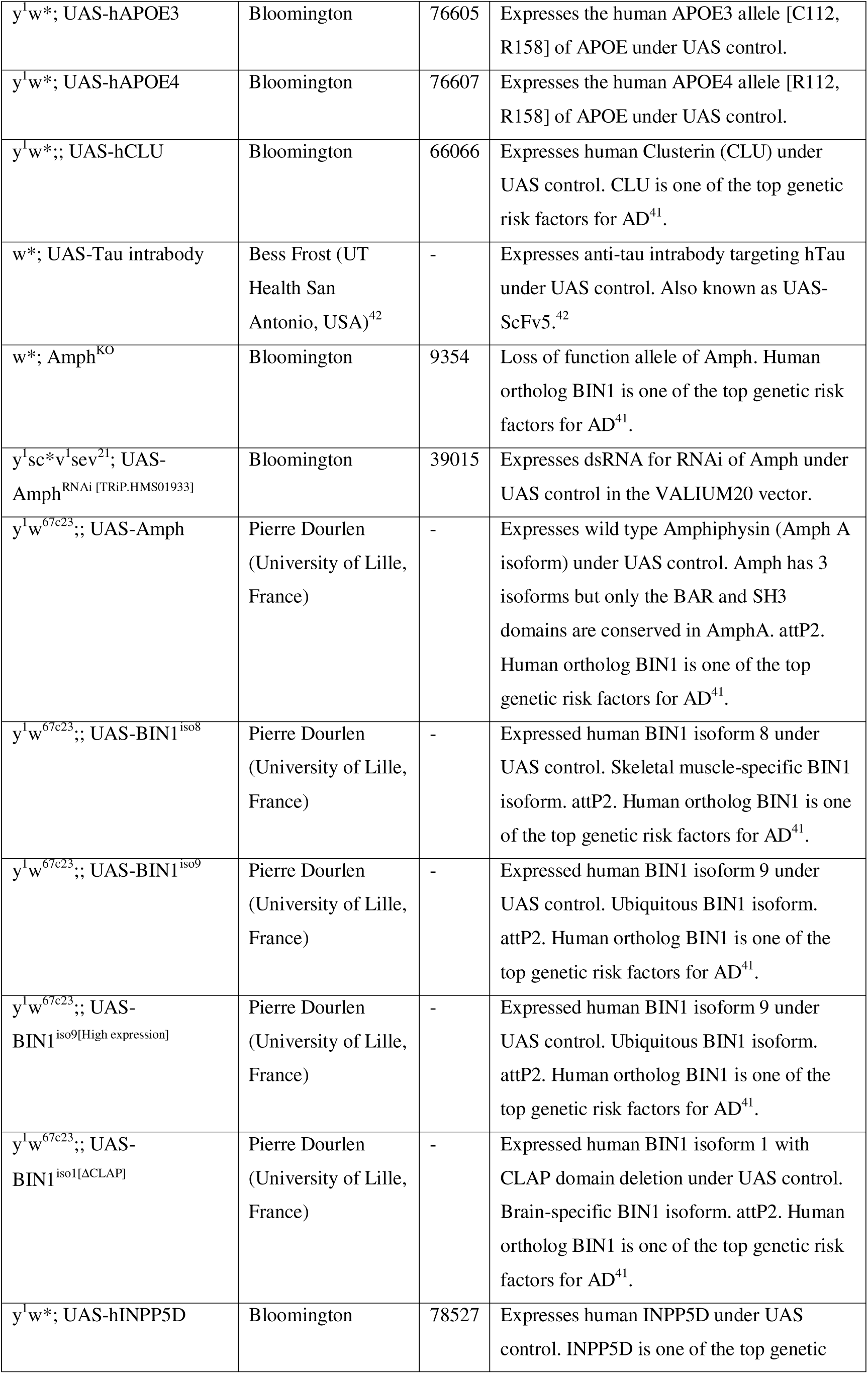
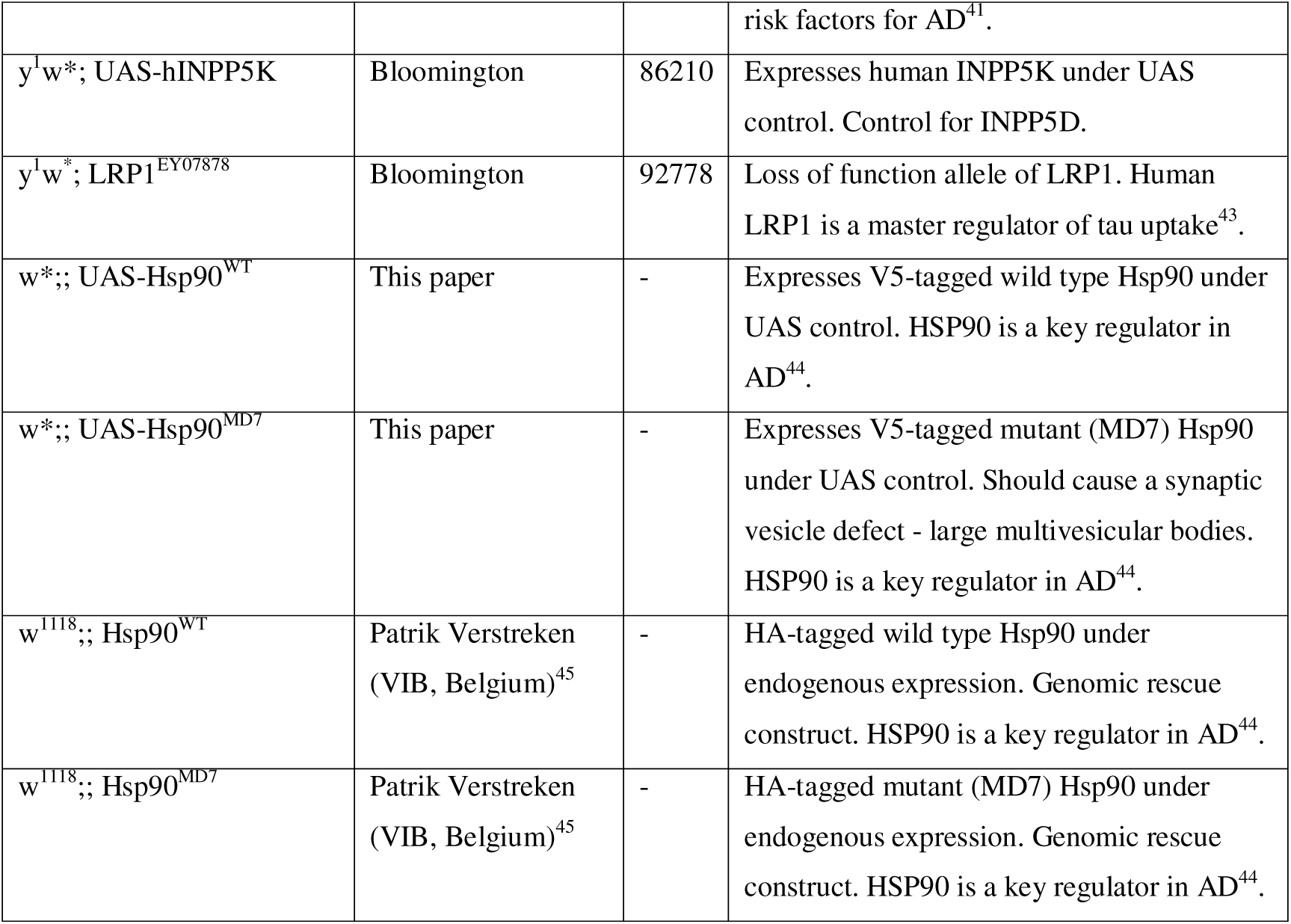
List of fly genotypes used.

### Generation of Repo>hAPOE2, Repo>hAPOE3, and Repo>hAPOE4 constructs

To generate the Repo>hAPOE2, Repo>hAPOE3, and Repo>hAPOE4 constructs, a new backbone plasmid (pRepo.attB) was generated by replacing the UAS promoter in the pUAST.attB plasmid with the *repo* promoter. The UAS promoter was removed from the plasmid by restriction digestion with HindIII and EcoRI.

The *repo* promoter region was amplified in two fragments from the bacterial artificial chromosome (BAC) clone CH321-23D05 using the following primers:

- FW_repo_p1: GAAGTTATGCTAGCGGATCCAACGAAACGAAAATGGGGAA
- RC_repo_p1: AAACTGGGAGGGATGGTAT
- FW_repo_p2: GGATACCATCCCTCCCAGTTT
- RC_repo_p2: GCGGCCGCAGATCTGTTAACGCTCGGCTACTGGCGATGAT

Both fragments were assembled into the linearized plasmid backbone using HiFi DNA Assembly (New England Biolabs), generating the intermediate vector pRepo.attB. Subsequently, the coding sequences (cDNA) of human APOE2, APOE3, and APOE4 were amplified from the following plasmids:

- pCMV4-ApoE2 (Addgene plasmid #87085)
- pCMV4-ApoE3 (Addgene plasmid #87086)
- pCMV4-ApoE4 (Addgene plasmid #87087)

Amplification was performed using the following primers:

- FW Kozak-ATG-v5-ApoE: AGCCGAGCGTTAACAGATCTGCAACATGGGTAAGCCAATTCCAAACCCATTGCTGGGTTTAGATTCTACTGGCGGTGGTGGTTCTAAGGTTCTGTGGGCTG CGT
- RC Kozak-ATG-v5-ApoE: AGGTTCCTTCACAAAGATCCTTCAGTGATTGTCGCTGGGC

The amplified APOE cDNA fragments contain a v5 epitope tag followed by a short linker (GGGS) and were cloned into the pRepo.attB vector following restriction digestion with NotI and XbaI, using HiFi DNA Assembly according to the manufacturer’s instructions. All final constructs were verified by Sanger sequencing prior to downstream applications.

### Generation of UAS-V5-Hsp90^WT^ and UAS-V5-Hsp90^MD7^ constructs

To generate the UAS-V5-Hsp90^WT^ and UAS-V5-Hsp90^MD7^ constructs, the coding sequences (cDNA) of Hsp90 ^WT^ and Hsp90 ^MD7^, together with the V5 epitope tag, were amplified from the following plasmids:

- pRS416-GAL1-v5-Hsp90[WT]^45^
- pRS416-GAL1-v5-Hsp90[MD7]^45^

Amplification was performed using the following primers:

- F-UAS-V5: aattcgttaacagatctgcggccgcggcAAAAAATGGGTAAGCCAATTCCAAA
- R-UAS-hsp90: ggttccttcacaaagatcctctagaTTAATCGACCTCCTCCATGTGGGAA

The amplified fragments were cloned into the pUAST.attB vector following restriction digestion with XhoI and XbaI, using HiFi DNA Assembly (New England Biolabs) according to the manufacturer’s instructions. All final constructs were verified by Sanger sequencing prior to downstream applications.

### Lifespan assay

Lifespan assays were performed at 25 °C under a 12 h:12 h light–dark cycle, adapted from established protocols^46^. Eggs were synchronised by collection on grape juice agar plates supplemented with yeast paste and approximately 300 eggs were transferred to each food bottle. Adults began eclosing from day 9; flies eclosing on the first day were discarded. One- day-old adults were allowed to mate for 48 h before being briefly anaesthetised with CO□, sorted by sex, and housed at 15 flies per vial (10 replicate vials per condition). Flies were transferred to fresh food three times per week and deaths were recorded at each transfer until all flies had died.

### Whole-mount brain immunostaining

Immunostaining was adapted from published whole-mount brain protocols with study-specific modifications^47^. Adult flies were immobilised with CO□ and fixed as whole animals prior to brain dissection. Up to 15 flies per condition were transferred to 1.5 mL tubes (Greiner bio-one 616 201) and fixed in 1.8 mL 4% paraformaldehyde (Agar Scientific, AGR1026) in PBS containing 0.5% Triton X-100 (PBS-T; Sigma X100-1L). Samples were incubated for 3 h at room temperature on a shaker with tubes positioned sideways to ensure full immersion. Flies were washed 4× for 15 min in 0.5% PBS-T. Brains were dissected in dilute detergent buffer (0.008% PBS-T) and transferred to 0.6 mL tubes. For phalloidin and DAPI counterstaining, brains were incubated for 16–24 h at 4°C in a cocktail containing DAPI (Sigma D9542; 1:1000) and Alexa Fluor phalloidin (Thermo Fisher A12381; 1:100) prepared in 0.5% PBS-T. Brains were washed 4× 15 min in 0.5% PBS-T and then 1× 30 min in 1× PBS to remove residual detergent.

#### Mounting

Brains were equilibrated in a 50:50 mixture of Vectashield mounting medium (Vector Laboratories H1200) and PBS-T prior to mounting. Brains were arranged on polylysine-coated microscope slides. To preserve tissue architecture, a bridge was created using two circular coverslips (VWR 631-0148) placed on either side of the tissue before applying a rectangular coverslip (VWR 631-0136). Coverslip edges were sealed with clear nail varnish and slides were stored protected from light until imaging.

#### Microscopy and image analysis for vacuolar degeneration and brain size

Image stacks spanning the entire brain were acquired on a Leica TCS SP8 multiphoton microscope using a 25× objective (Coherent Chameleon laser, 760 nm; LAS X software). Images were processed in Fiji/ImageJ. Files were blinded prior to quantification using the Blind Analysis Tools plug-in^48^. For brain area, the z-plane with the largest cross-sectional area (typically mid-stack) was identified by manual inspection; the brain outline was traced using the polygon selection tool and area recorded. Vacuoles were quantified in the optic lobes using the Cell Counter plug-in. Optic lobes were analysed due to their consistent and readily quantifiable degeneration across individuals and genotypes. Vacuoles were counted through the z-stack with care to count each vacuole once even if spanning adjacent planes.

### Western blotting

#### Sample preparation

Fly heads were lysed directly in SDS sample buffer on ice. For each head, 10 µL of lysis buffer was prepared consisting of 5 µL 2× SDS sample buffer (Sigma-Aldrich S3401-1VL), 0.5 µL 1 M DTT (final 50 mM; Sigma-Aldrich), and 4.5 µL dH□O. Samples typically comprised 15 heads per replicate (150 µL total volume), with four biological replicates per group.

#### Gel electrophoresis and transfer

Samples were heated at 80°C for 10 min, vortexed, and centrifuged at maximum speed for 2 min. Proteins were resolved on 4–12% Bis-Tris gels (NuPAGE; Thermo Fisher; 15-well format) in MES running buffer (NuPAGE MES SDS 20×, Invitrogen NP0002). Each lane received 10 µL sample; molecular weight marker (LI-COR 928-40000) was loaded at 5 µL. Electrophoresis was performed at 80 V for 5 min followed by 120 V for 1.5 h. Gels were equilibrated in 20% ethanol for 10 min prior to transfer. Proteins were transferred to PVDF membranes (included with the iBlot3 stack) using the iBlot3 semi-dry transfer system (Invitrogen IB31000) with iBlot3 PVDF transfer stacks (mini format) according to manufacturer instructions (8.5 min at 20 V, no cooling), ensuring bubble-free stack assembly.

#### Total protein staining, immunodetection, and imaging

Membranes were stained using REVERT 700 Total Protein Stain (LI-COR 926-11021) and imaged on an Odyssey Fc Imager (700 nm channel). Membranes were subsequently destained using REVERT Destaining Solution (LI-COR 926-11023; 8 min) before immunoblotting.

Membranes were blocked in Intercept Blocking Buffer (LI-COR 927-70001) for 60 min and incubated overnight in primary antibodies diluted in Intercept buffer with 0.1% Tween-20. Primary antibodies included mouse anti-Tau5A6 (DSHB, 1:500), rabbit anti-synaptogyrin^39^ (gift from T. Littleton; 1:10,000) and mouse anti-Dlg1 (DSHB; 1:1,000). After washing (6× 5 min, PBS), membranes were incubated with IRDye secondary antibodies (1:5,000) for 1.5 h at room temperature protected from light (IRDye 800CW donkey anti-rabbit, LI-COR 925-32213; IRDye 680RD donkey anti-mouse, LI-COR 925-68072). Membranes were washed (6× 5 min, PBS) and imaged on the Odyssey Fc system. Quantification was performed in Empiria Studio (LI-COR) using guided analysis workflows.

### *Drosophila* activity monitoring and sleep analysis

Locomotor activity was recorded using 32-channel DAM2 monitors (DAM2, TriKinetics) with 5 mm × 65 mm polycarbonate tubes (PPT5×65, TriKinetics) sealed with vinyl caps (CAP5-Black, TriKinetics); air-permeable plugs were prepared from diced cellulose acetate plugs (Droso-Plugs, Genesee 59-200). Sleep was defined operationally as a period of at least five minutes of inactivity, as detected by the absence of beam crossings. This threshold reflects key features of sleep in *Drosophila*, such as its circadian regulation, recovery after deprivation, diminished sensory responsiveness during extended inactivity, characteristic body posture, and brain state changes occurring over similar time intervals^49–52^. While sleep inferred through DAM is based on inactivity and may not separate sleep from quiet wakefulness, it has been shown to be a reliable and scalable behavioural proxy for assessing sleep in flies^52^.

DAM food was prepared by dissolving agar (1%, 2.5 g; Sigma-Aldrich, A7002-100G) and sugar (5%, 13 g; Sigma-Aldrich, S9378-1KG) in 250 mL distilled water. The solution was heated to boiling to ensure complete dissolution and then cooled to a workable temperature. For standard RU induction, RU486 was added to a final concentration of 200 µM. For dose-response experiments, RU486 was added at the indicated concentrations: 10, 50, 100, or 200 µM. Tubes were filled by standing upright in molten medium until solidified. Monitors were housed in an incubator at 25°C under 12 h:12 h LD cycles.

Flies were briefly anaesthetised with CO□ and loaded individually into tubes, plugged, and placed horizontally until recovery. Flies were then mounted in DAM units and recorded for 6 days. The first full day under LD was excluded to allow acclimation. Data from three consecutive LD cycles (days 3–5) were extracted for analysis.

Raw monitor files were processed using the standalone ShinyR-DAM application^12^. The application generated derived measures including sleep (defined as ≥5 consecutive minutes of inactivity), activity and sleep profiles, and bout metrics, and exported parameters as .csv files. These files were then analysed in custom R scripts adapted from the ShinyR-DAM supplementary analysis pipeline^12^.

### Statistical analysis

Statistical analyses and data visualisation were performed in R/RStudio^53^ and Microsoft Excel using the Piper Lab lifespan template^54^. Full analysis workflows, including data processing, model diagnostics, sensitivity analyses, and code, were generated as R Markdown/HTML reports and will be made available with the associated data. Primary models were selected according to the structure of each response variable and the experimental design. The main statistical approaches are summarised in **Table 2**.

**Table 2.**
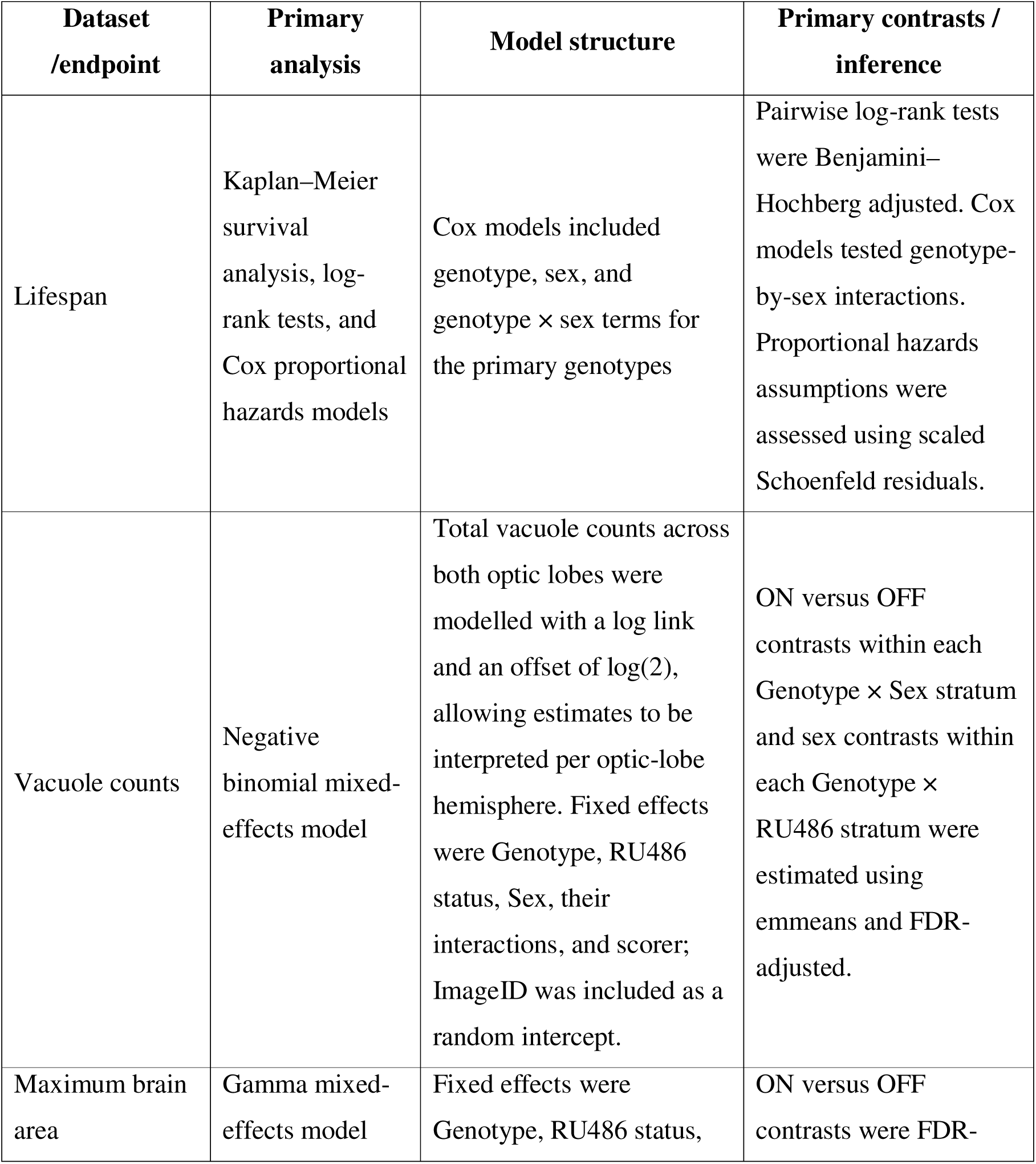
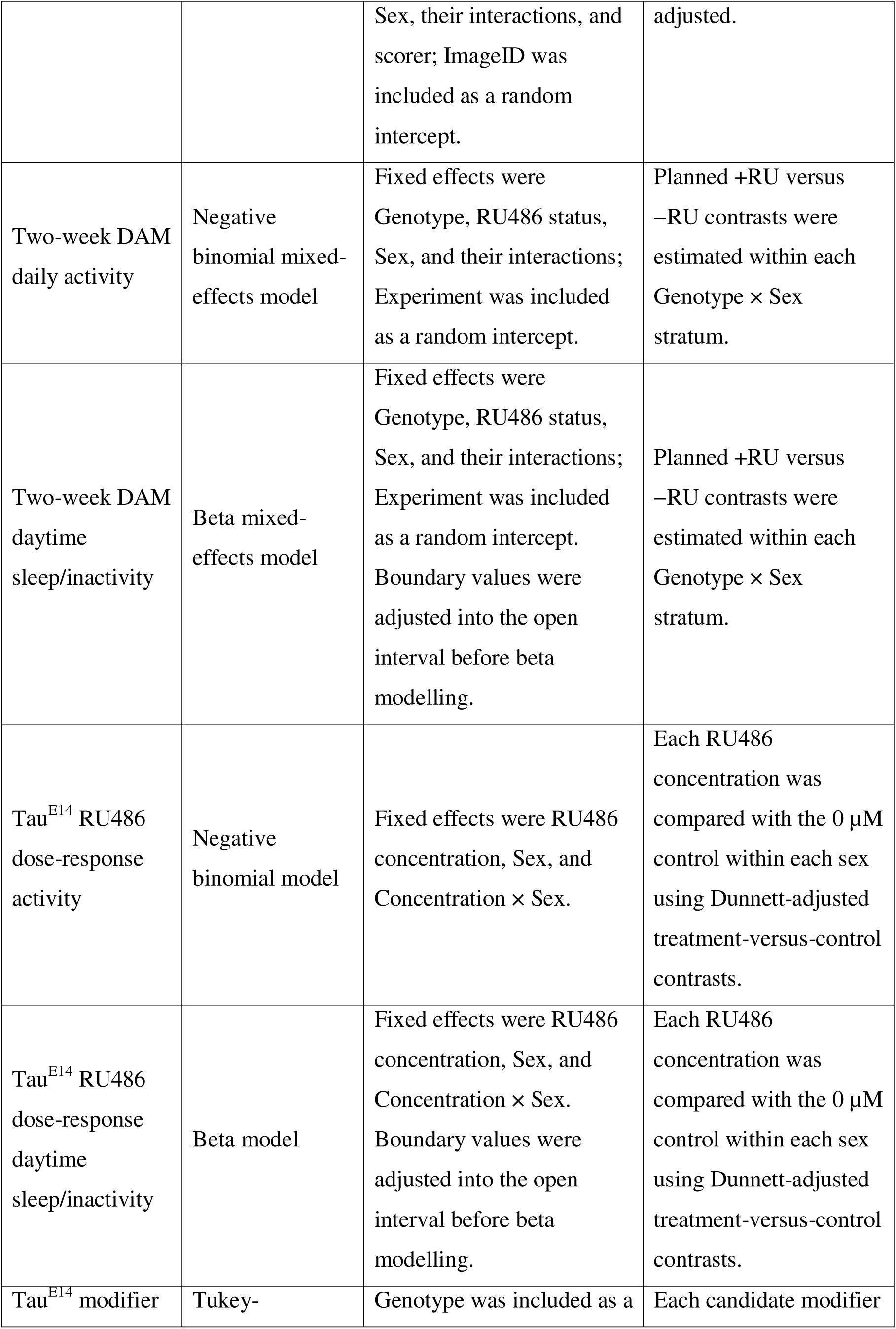
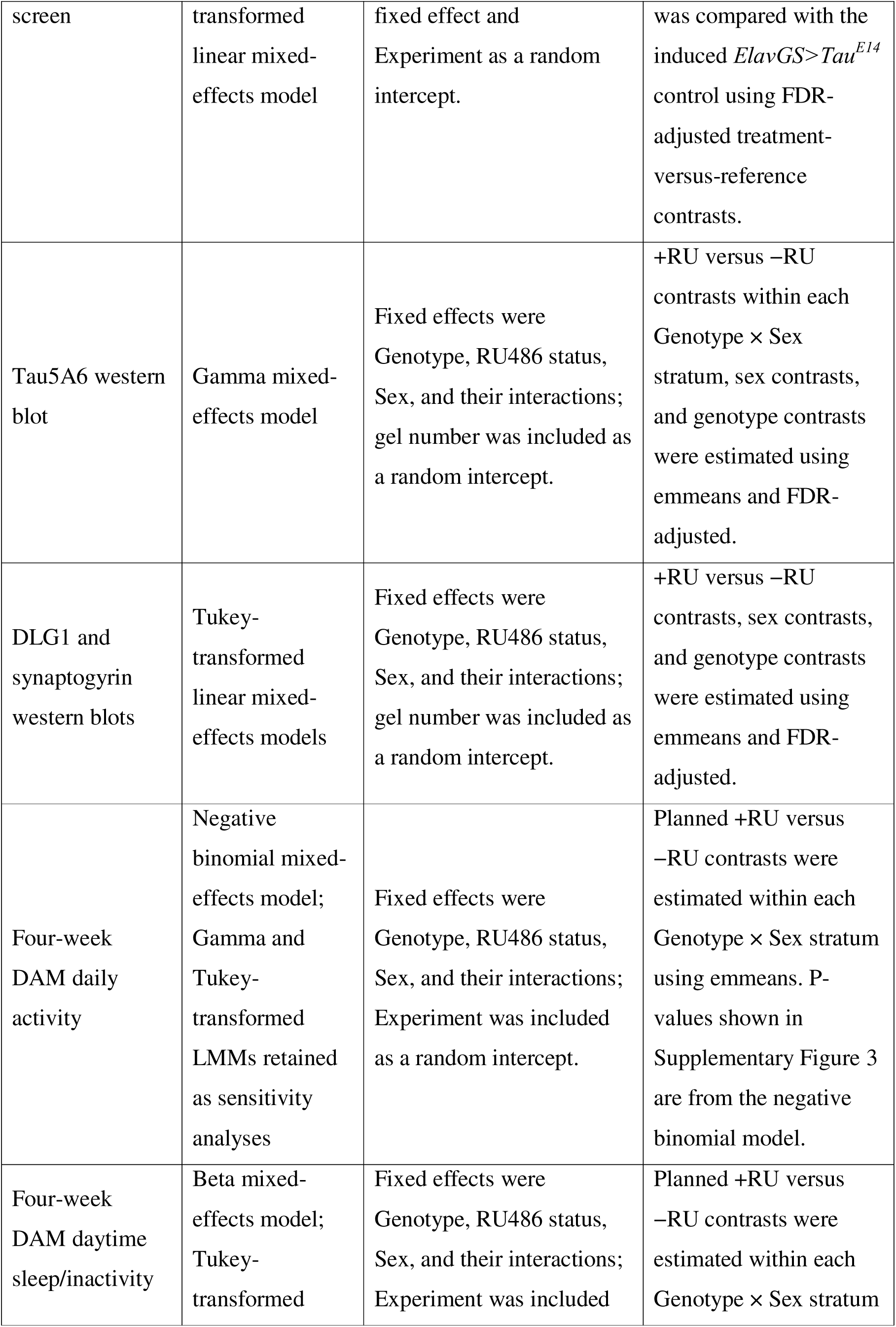
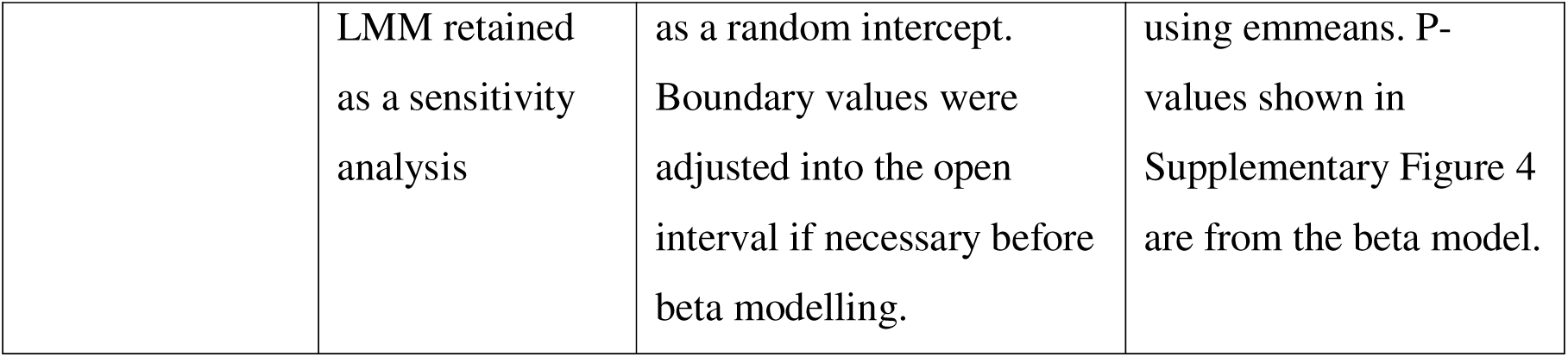
Summary of primary statistical analyses.

For mixed-effects models, Type III tests were used to assess fixed effects. For generalized mixed models, Type III Wald χ^2^ tests were used. Planned contrasts were estimated using emmeans. Multiple-testing correction was applied as described for each analysis, using false discovery rate correction unless otherwise specified. Model diagnostics included inspection of residual distributions and variance patterns, simulation-based diagnostics where appropriate, and assessment of model assumptions using performance, DHARMa, and ggResidpanel. Sensitivity analyses using alternative response transformations or distributional assumptions were performed for key endpoints and are reported in the accompanying statistical analysis files. Statistical significance was defined at α = 0.05.

### Software

Figures and schematics were generated using BioRender and Inkscape. Behavioural and statistical analyses were performed in RStudio. Western blot acquisition and quantification were performed using Image Studio and Empiria Studio (LI-COR). Vacuolar images were analysed and quantified using FIJI/ImageJ. Cartoons were generated in BioRender. Artificial intelligence tools (ChatGPT, OpenAI) were used to assist with R programming and language editing during manuscript preparation. All analyses, interpretation of results, and final manuscript content were reviewed and verified by the authors.

## Results

### Adult-onset pan-neuronal tau expression reduces lifespan and induces neurodegeneration

Because pan-neuronal expression of human tau during development causes severe deleterious neurodevelopmental abnormalities in *Drosophila* (including abnormal eye development), we asked whether restricting tau expression to adulthood is sufficient to drive neurodegeneration in a fashion more similar to human late onset tauopathies than developmental phenotypes.

To test whether adult-onset neuronal tau expression is sufficient to reduce survival, we induced pan-neuronal expression of wild-type 0N4R tau (Tau^WT^) or the phospho-mimetic variant Tau^E14^ beginning in adulthood using the RU486-inducible *ElavGS* driver. Survival was assessed separately in females and males, alongside driver-alone and UAS-alone controls. In both sexes, adult-onset expression of either Tau^WT^ or Tau^E14^ resulted in a pronounced reduction in lifespan relative to control genotypes (**Figure 1A,B**). Median lifespan in *ElavGS* controls was 63 days in females and 60 days in males, compared with 46 and 53 days in Tau^WT^-expressing females and males, and 35 and 37 days in Tau^E14^-expressing females and males, respectively. Pairwise Benjamini–Hochberg-adjusted log-rank tests showed that tau-expressing flies had significantly reduced survival relative to control genotypes in both sexes, with Tau^E14^ producing the strongest effect. When normalised to sex-matched *ElavGS* controls, Tau^WT^ reduced median lifespan by ∼27% in females and ∼12% in males, whereas Tau^E14^ reduced median lifespan by ∼44% and ∼38% in females and males, respectively.

**Figure 1.**
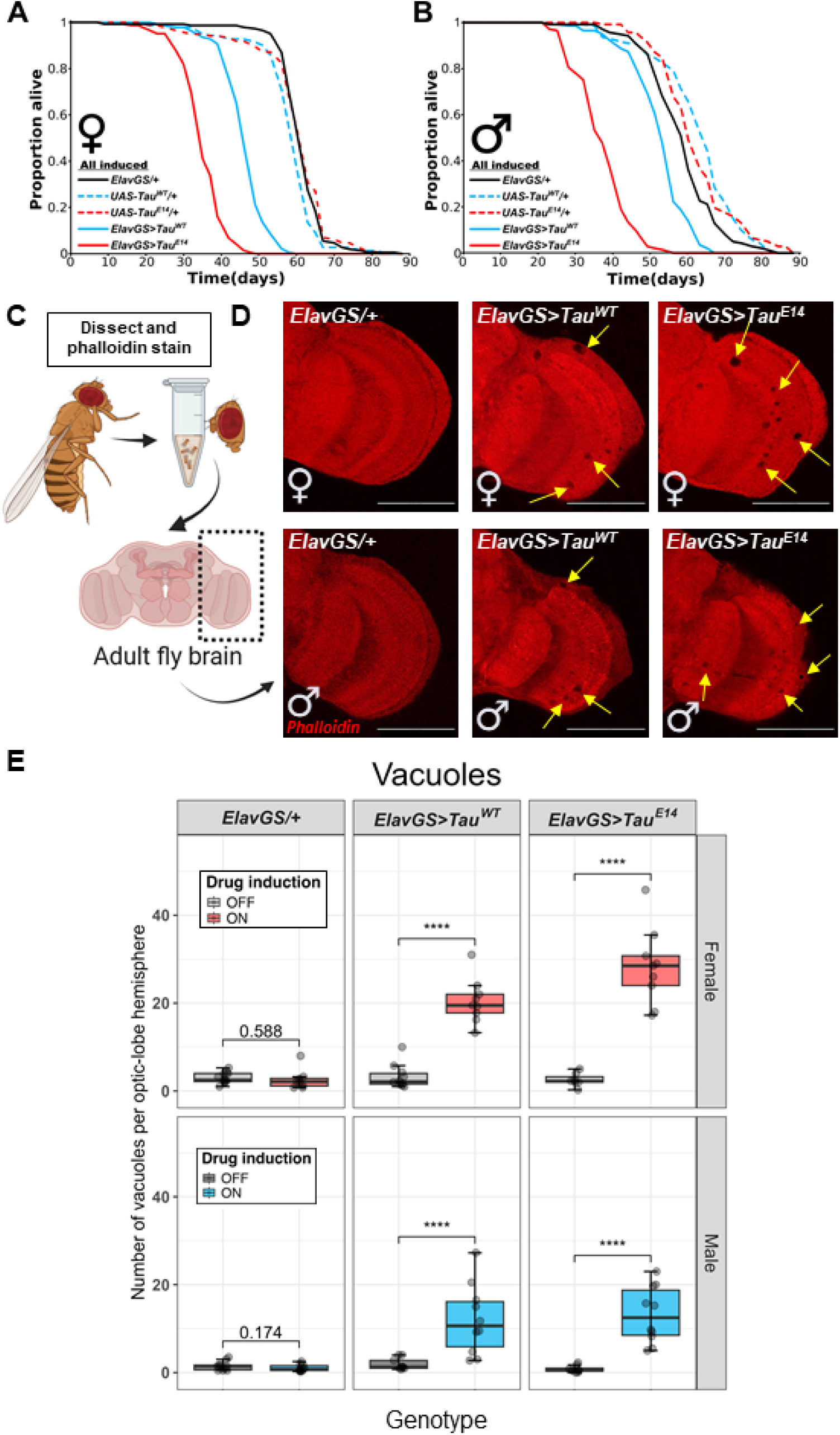
Adult-onset pan-neuronal tau expression reduces lifespan and induces neurodegeneration. (**A**) Female survival curves for control flies (*ElavGS/+* and *UAS-Tau lines*) and flies expressing human Tau pan-neuronally (*ElavGS>Tau^WT^* and *ElavGS>Tau^E^*^14^). (**B**) Male survival curves for the same genotypes. Flies were raised at 25 °C and maintained on RU486-containing food to induce adult-onset expression of Tau. n = 142–151 flies per condition; censored flies were included in survival analyses. In both sexes, lifespan was shortest in *ElavGS>Tau^E^*^14^ flies, followed by *ElavGS>Tau^WT^*, compared to control genotypes. (**C**) Schematic illustrating dissection of the adult fly brain and the orientation of the optic lobes imaged for analysis. (**D**) Representative micrographs of optic lobes stained with phalloidin to visualise neuropil structure in induced flies (all fed RU food to induce expression in adult neurons). Expression of Tau leads to the formation of vacuolar lesions characteristic of neurodegeneration. Scale bar: 100 µm. (**E**) Quantification of vacuole number in the optic lobes after four weeks of tau induction. Vacuoles in the left and right optic lobes were manually counted by two blinded scorers. Points show individual flies/brains, with vacuole counts averaged across scorers for visualisation (*n* = 7–11 per genotype). Statistical analysis was performed on scorer-level data using a negative binomial mixed-effects model fitted to total vacuole counts across both optic-lobe hemispheres, with an offset to express estimates per hemisphere. The model included Genotype, RU486 status, Sex, and their interactions as fixed effects, scorer as a fixed effect, and ImageID as a random intercept. Planned ON versus OFF contrasts were FDR-adjusted. Significance thresholds are indicated as *P* < 0.05 (\**), P < 0.01 (**), and P < 0.001 (\*\*\**). Cartoon created in BioRender. Tulloch, J. (2026) https://BioRender.com/adonv0q

To formally test whether tau effects differed by sex, we fitted a Cox proportional hazards model to the primary genotypes, including genotype, sex, and their interaction. This analysis revealed a significant genotype-by-sex interaction likelihood-ratio test: χ^2^(2) = 58.18, p = 2.3 × 10^-13, indicating that the magnitude of tau-induced lifespan shortening differed between males and females. Sex-specific Cox models further supported stronger tau-associated mortality effects in females than males. Compared with *ElavGS* controls, Tau^WT^ expression increased mortality hazard more strongly in females than males, and the same pattern was observed for Tau^E14^. Thus, adult-onset tau expression reduces lifespan in both sexes, but the effect is more severe in females. Given this robust reduction in lifespan, we next examined whether adult-onset, pan-neuronal tau expression was associated with overt neurodegeneration.

Vacuole formation was quantified in the optic lobes after four weeks of RU486 induction, a time point selected to precede the lowest median lifespan observed in survival assays. Although vacuolisation was observed in multiple brain regions, the optic lobes were analysed because degeneration in this area was consistent and readily quantifiable across individuals and genotypes (**Figure 1C** and **Supplementary Figure 1A**). Marked vacuole formation was observed in the optic lobes of Tau^WT^- and Tau^E14^-expressing flies under induction (+RU), whereas uninduced controls (−RU) showed minimal degeneration (**Figure 1D,E** and **Supplementary Figure 1A**).

Vacuole counts were analysed using a negative binomial mixed-effects model fitted to scorer-level data, with ImageID included as a random intercept and scorer included as a fixed effect. This analysis revealed significant effects of Genotype (χ^2^(2) = 87.89, p < 0.0001), RU486 status (χ^2^(1) = 151.73, p < 0.0001), Sex (χ^2^(1) = 48.16, p < 0.0001), and a strong Genotype × RU486 status interaction (χ^2^(2) = 109.48, p < 0.0001). FDR-adjusted contrasts showed that RU486 induction significantly increased vacuole burden in both Tau^WT^- and Tau^E14^-expressing flies in females and males (all p < 0.0001), but not in *ElavGS/+* controls (female p = 0.588; male p = 0.174). On the response scale, induction increased estimated vacuole burden by approximately 7.2-fold in Tau^WT^ females, 6.2-fold in Tau^WT^ males, 13.1-fold in Tau^E14^ females, and 16.9-fold in Tau^E14^ males (**Figure 1E**).

Although there was a significant main effect of sex, with males generally showing lower vacuole counts than females, there was no significant Genotype × Sex interaction (χ^2^(2) = 3.33, p = 0.189), RU486 status × Sex interaction (χ^2^(1) = 0.051, p = 0.821), or Genotype × RU486 status × Sex interaction (χ^2^(2) = 0.83, p = 0.661). Thus, while overall vacuole burden differed between sexes, the induction-dependent tau-associated increase in vacuolisation was broadly similar in males and females at this structural level (**Supplementary Figure 1B**).

RU induction increased maximum cross-sectional brain area in females but had little effect in males. Consistent with this, analysis using a Gamma mixed-effects model revealed significant effects of RU486 status (χ^2^(1) = 53.36, p = 2.8 × 10□¹³) and sex (χ^2^(1) = 24.18, p = 8.8 × 10□□), as well as a strong RU486 status × Sex interaction (χ^2^(1) = 53.71, p = 2.3 × 10□¹³) (**Supplementary Figure 1C**). FDR-adjusted contrasts showed that RU486 increased brain area in females across genotypes (all p < 0.0001), but not in males (all p ≥ 0.262), indicating that this effect was sex-dependent and not specific to tau expression.

In total, these findings show that adult-onset pan-neuronal tau expression is sufficient to reduce lifespan and induce reproducible structural neurodegeneration in the *Drosophila* brain. While lifespan analyses revealed clear sex-dependent modulation of tau toxicity, the vacuole analysis did not reveal a strong sex-specific modulation of the induction-dependent tau effect, suggesting that sex differences in tau toxicity may arise through mechanisms not fully captured by gross vacuolar neurodegeneration alone. Vacuolar pathology was detectable well before median lifespan was reached, suggesting that degeneration develops progressively following tau induction. We therefore used automated DAM-based behavioural assays to test whether earlier functional changes could be detected.

### Adult-onset tau expression induces sexually dimorphic changes in locomotor activity during early adulthood

To examine behavioural consequences of adult-onset tau expression, locomotor activity and daytime sleep/inactivity were analysed after two weeks of induction using the DAM system under 12 h:12 h light–dark conditions. Daily locomotor activity was analysed using a negative binomial mixed-effects model, while daytime sleep/inactivity, a bounded proportional measure, was analysed using a beta mixed-effects model.

Tau expression produced a striking sex-dependent behavioural phenotype. For daily locomotor activity, the negative binomial model revealed a significant Genotype × RU486 status × Sex interaction (χ^2^(4) = 78.67, p < 0.0001). In males, induction of either Tau^WT^ or Tau^E14^ significantly increased daily activity relative to genotype-matched uninduced controls (both p < 0.0001). In females, Tau^E14^ induction significantly reduced activity (p = 0.0055), whereas Tau^WT^ induction did not significantly alter total daily activity (p = 0.8318) (**Supplementary Figure 2**). RU486 also increased activity in *ElavGS/+* driver controls in both females and males (both p < 0.0001), indicating that RU486 exposure itself can influence locomotor output.

Daytime sleep/inactivity showed a stronger and more consistent tau-dependent sex divergence (Figure 2A**,B**). Beta mixed-effects modelling revealed a significant Genotype × RU486 status × Sex interaction (χ^2^(4) = 150.31, p < 0.0001). In females, induction of either Tau^WT^ or Tau^E14^ significantly increased daytime sleep/inactivity relative to uninduced genotype-matched controls (both p < 0.0001). In contrast, males expressing Tau^WT^ or Tau^E14^ showed significantly reduced daytime sleep/inactivity upon induction (both p < 0.0001).

**Figure 2.**
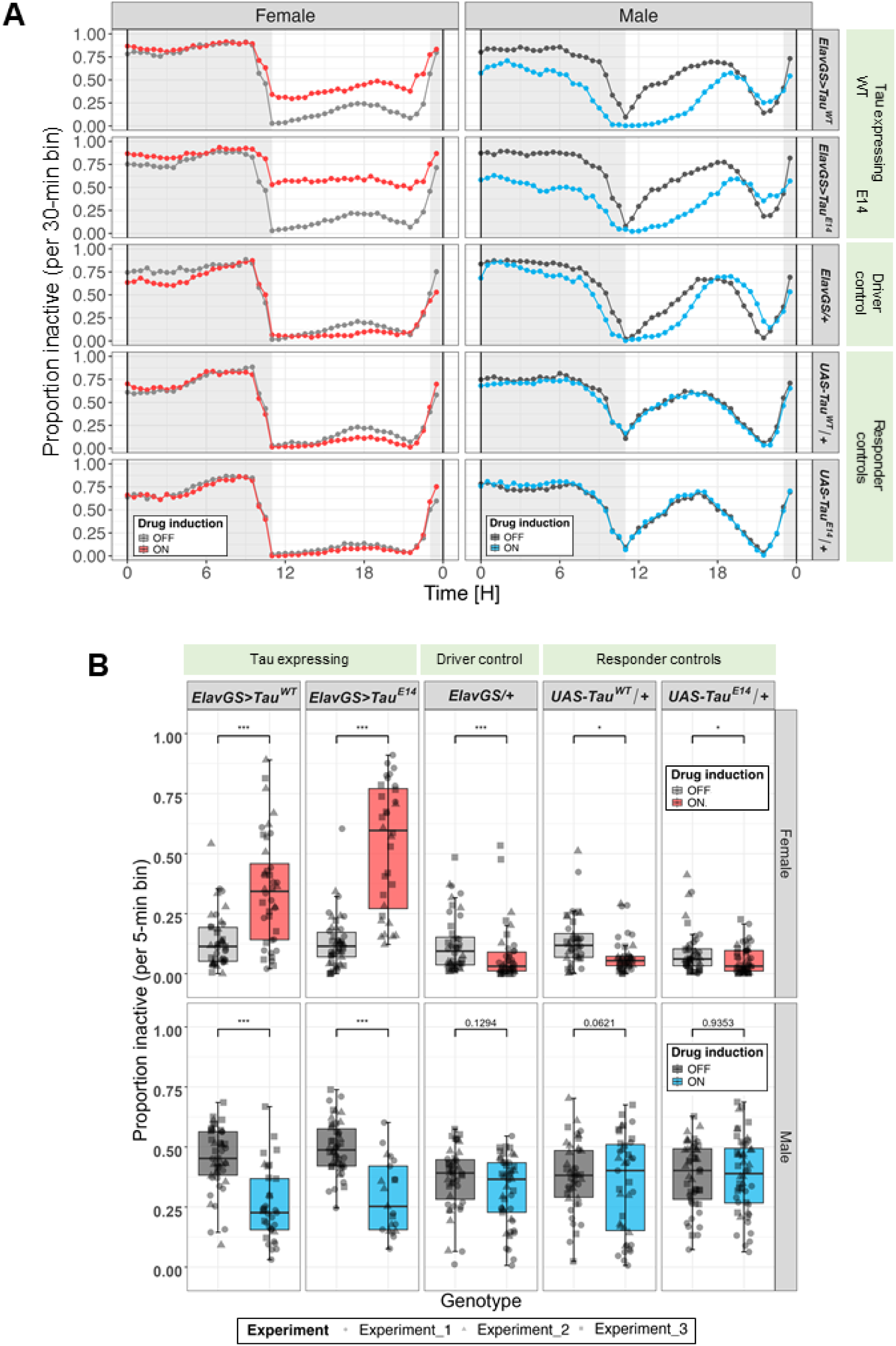
Adult-onset pan-neuronal expression of human Tau for 2 weeks alters daytime inactivity in a sexually dimorphic manner. (**A**) Average daily inactivity profiles (3-day mean) under 12:12 light–dark conditions showing population mean inactivity across the day. Light hours (day) are shown with a white background and dark (night) hours shown with a grey background. Inactivity values represent the proportion of inactive 5-minute bins averaged within each 30-minute interval. Female Tau^WT^ and Tau^E14^ expressing flies show increased daytime inactivity compared to controls, whereas male Tau^WT^ and Tau^E14^ expressing flies show reduced inactivity. Profiles represent data averaged from three independent experiments. Flies were 2 weeks old when introduced into the DAMS. (**B**) Boxplots showing mean daytime inactivity for each genotype. Adult-onset pan-neuronal expression of Tau^WT^ or Tau^E14^ resulted in increased daytime inactivity in females and decreased inactivity in males relative to controls. Individual points represent single flies and point shapes indicate independent experiments. Boxplot colours denote RU486 drug status: grey (–RU; no drug, Tau expression OFF) and coloured (+RU; drug administered, Tau expression ON; red = females, blue = males). Statistical analysis of daytime sleep/inactivity was performed using a beta mixed-effects model with Genotype, RU486 status, Sex, and their interactions as fixed effects, and Experiment as a random intercept. Pairwise comparisons show planned +RU versus −RU contrasts within each Genotype × Sex stratum. P-values are derived from the beta model. Significance thresholds are indicated as *P* < 0.05 (\**), P < 0.01 (**), and P < 0.001 (\*\*\**). Sample sizes ranged from *n* = 21–48 flies per condition.

Importantly, these tau-dependent effects were distinct from RU486 effects in control genotypes. In females, RU486 reduced daytime sleep/inactivity in *ElavGS/+* and UAS responder controls, opposite in direction to the increase observed with tau expression. In males, RU486 did not significantly alter daytime sleep/inactivity in control genotypes. Thus, the sexually dimorphic tau phenotype cannot be explained by RU486 exposure alone.

Together, these data show that two weeks of adult-onset tau expression is sufficient to produce robust behavioural disruption, with females exhibiting increased daytime sleep/inactivity and males showing reduced daytime sleep/inactivity and hyperactivity.

### Adult-onset tau expression produces dose-dependent, sexually dimorphic activity changes

To determine whether the behavioural phenotypes observed at two weeks scaled with the level of Tau^E14^ induction, we examined locomotor activity across a range of RU486 concentrations (0–200 μM). This dose-response analysis allowed us to assess whether the activity changes progressed gradually with increasing induction and whether the apparent sexual dimorphism might reflect differences in effective Tau dosage between sexes.

Daily locomotor activity was analysed using a negative binomial model with Concentration, Sex, and their interaction as predictors. This revealed a significant effect of concentration and a strong Concentration × Sex interaction, indicating that the dose-response relationship differed between females and males (Concentration: χ^2^(4) = 12.13, p = 0.0164; Sex: χ^2^(1) = 145.15, p < 0.0001; Concentration × Sex: χ^2^(4) = 153.07, p < 0.0001). Across the dose range, Tau^E14^ expression produced opposing behavioural responses in the two sexes (Figure 3). Activity profiles revealed clear dose-dependent changes in locomotor output (**Figure 3A,B**). Dunnett-adjusted treatment-vs-control contrasts showed opposing dose-dependent effects in the two sexes. In males, RU486 increased activity at all concentrations tested relative to 0 µM controls, including 10 µM (p = 0.0037), 50 µM (p = 0.0003), 100 µM (p < 0.0001), and 200 µM (p < 0.0001). In contrast, females showed reduced activity at 50, 100, and 200 µM (all p < 0.0001), with no significant change at 10 µM (p = 0.9974).

**Figure 3.**
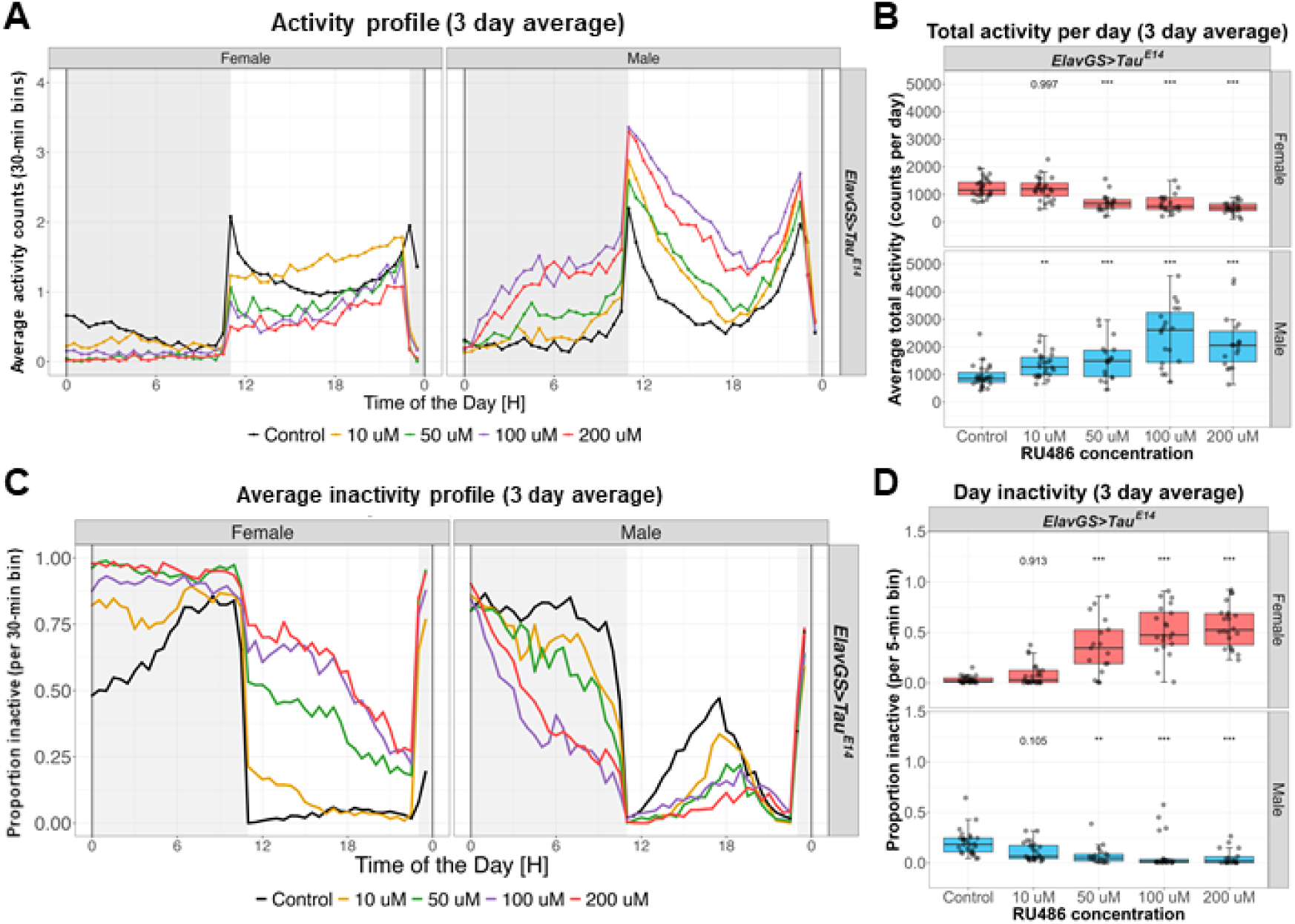
Dose-dependent effects of adult-onset Tau^E14^ expression on locomotor activity and daytime inactivity in *Drosophila*. (**A**) Average locomotor activity profiles (3-day mean) under 12:12 light–dark conditions for male and female *ElavGS>Tau^E^*^14^ flies exposed to increasing RU486 concentrations (Control, 10 µM, 50 µM, 100 µM, and 200 µM). Activity counts per minute are plotted across the 24-hour cycle. Activity counts were averaged in 30-minute bins for each genotype. Each line represents the mean locomotor activity for a given RU486 concentration. Shaded regions indicate the dark phase. (**B**) Boxplots showing average daily locomotor activity across RU486 concentrations. Increasing RU486 concentrations resulted in dose-dependent changes in activity in male and female flies expressing Tau^E14^. Individual points represent single flies and colours indicate sex (red = females, blue = males). (**C**) Average daytime inactivity profiles (3-day mean) under 12:12 light–dark conditions for the same genotypes and RU486 concentrations. Inactivity values represent the proportion of inactive time across the day. Each line represents the mean inactivity profile for a given RU486 concentration. (**D**) Boxplots showing mean daytime inactivity across RU486 concentrations. Individual points represent single flies and colours indicate sex (red = females, blue = males). Flies were 2-weeks old when introduced into the DAMS. Daily locomotor activity was analysed using a negative binomial model with Concentration, Sex, and their interaction as predictors. Daytime sleep/inactivity was analysed using a beta model with the same predictor structure. Planned treatment-vs-control contrasts compared each RU486 concentration with the 0 µM control within each sex and were Dunnett-adjusted. Significance thresholds are indicated as *P* < 0.05 (\**), P < 0.01 (**), and P < 0.001 (\*\*\**). N = 17-27 females and 18-27 males.

Consistent with these activity changes, inactivity profiles showed a reciprocal pattern (**Figure 3C,D**). Daytime sleep/inactivity was analysed using a beta model because the response is a bounded proportional measure. This analysis showed significant effects of Concentration and Sex, together with a strong Concentration × Sex interaction (Concentration: χ^2^(4) = 32.01, p < 0.0001; Sex: χ^2^(1) = 53.26, p < 0.0001; Concentration × Sex: χ^2^(4) = 154.07, p < 0.0001). Consistent with the activity data, daytime sleep/inactivity showed reciprocal dose-dependent changes. In females, daytime sleep/inactivity was significantly increased at 50, 100, and 200 µM relative to 0 µM controls (all p < 0.0001), with no significant change at 10 µM (p = 0.9129). In males, daytime sleep/inactivity was significantly reduced at 50 µM (p = 0.0039), 100 µM (p = 0.0001), and 200 µM (p < 0.0001), while the reduction at 10 µM did not reach significance (p = 0.1049). Together, these analyses indicate that tau induction produces a dose-dependent and sexually dimorphic behavioural response at the two-week time point.

In the four-week activity dataset, negative binomial mixed-effects modelling revealed a significant Genotype × RU486 status × Sex interaction (χ^2^(4) = 48.18, p < 0.0001). RU486 induction reduced activity in Tau^WT^ females and Tau^E14^ females, increased activity in Tau^WT^ males, but did not significantly alter total daily activity in Tau^E14^ males. Daytime inactivity analysed using a beta mixed-effects model also showed a significant Genotype × RU486 status × Sex interaction (χ^2^(4) = 80.88, p < 0.0001); induction increased daytime inactivity in Tau^WT^ and Tau^E14^ females and in Tau^E14^ males, but not in Tau^WT^ males.

### Adult-onset tau expression is robustly induced and is not accompanied by loss of DLG1 or synaptogyrin after two weeks

To confirm effective induction of the GeneSwitch system, total human tau levels were assessed in whole-head lysates after two weeks of adult-onset expression in *ElavGS>Tau^WT^*and *ElavGS>Tau^E^*^14^ flies, alongside *ElavGS/+* controls. Western blotting using the Tau5A6 antibody confirmed robust RU486-dependent induction of human tau in both tau-expressing genotypes (**Figure 4A,B; Supplementary Figure 5**). Tau5A6 signal was analysed using a Gamma mixed-effects model with Genotype, RU486 status, Sex, and their interactions as fixed effects, and gel number as a random intercept. This revealed strong effects of Genotype (χ^2^(2) = 467.95, p < 0.0001), RU486 status (χ^2^(1) = 200.70, p < 0.0001), and a Genotype × RU486 status interaction (χ^2^(2) = 83.59, p < 0.0001), consistent with selective induction of tau in *ElavGS>Tau^WT^* and *ElavGS>Tau^E^*^14^ flies. Planned +RU versus −RU contrasts showed significant induction in Tau^WT^ and Tau^E14^ flies in both females and males (all p < 0.0001), but not in *ElavGS/+* controls.

**Figure 4.**
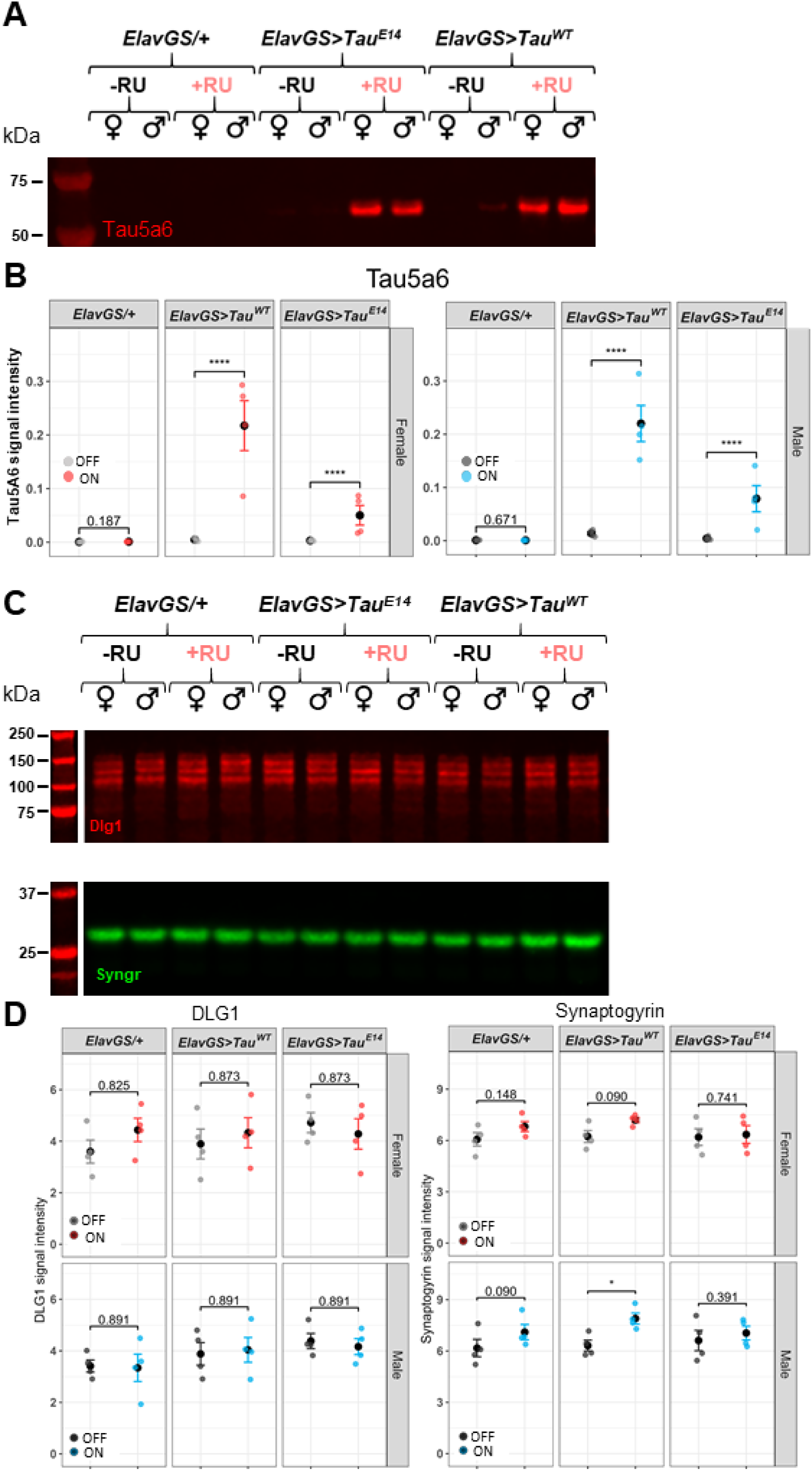
Adult-onset tau expression is robustly induced and is not accompanied by loss of DLG1 or synaptogyrin after two weeks. (**A**) Representative western blot showing Tau5A6 signal in whole-head lysates from male and female *ElavGS/+, ElavGS>Tau^WT^*, and *ElavGS>Tau^E^*^14^ flies after two weeks on control food or RU486-containing food. (**B**) Quantification of Tau5A6 signal normalised to total protein. Individual points represent biological replicates from independent gels (n = 4 per condition). RU486 induced robust Tau5A6 signal in *ElavGS>Tau^WT^* and *ElavGS>Tau^E^*^14^ flies, but not in *ElavGS/+* controls. Statistical analysis was performed using a Gamma mixed-effects model with Genotype, RU486 status, Sex, and their interactions as fixed effects, and gel number as a random intercept. Planned +RU versus −RU contrasts within each Genotype × Sex stratum were FDR-adjusted. (**C)** Representative western blots showing DLG1 and synaptogyrin in whole-head lysates from the indicated genotypes after two weeks of adult-onset tau expression. (**D**) DLG1 and synaptogyrin levels were quantified from whole-head lysates and normalised to total protein. Individual points represent biological replicates from independent gels (n = 4 per condition). Statistical analysis was performed using Tukey-transformed linear mixed-effects models with Genotype, RU486 status, Sex, and their interactions as fixed effects, and gel number as a random intercept. Planned +RU versus −RU contrasts within each Genotype × Sex stratum were FDR-adjusted. No induction-dependent loss of DLG1 or synaptogyrin was detected in tau-expressing flies. Individual points represent biological replicates. Significance thresholds are indicated as P < 0.05 (*), P < 0.01 (**), and P < 0.001 (***).

Low Tau5A6 signal was detectable in some uninduced tau-transgene samples, consistent with low basal signal or leakiness, but RU486 induction increased Tau5A6 signal by approximately 48-fold in Tau^WT^ females, 17-fold in Tau^WT^ males, 17-fold in Tau^E14^ females, and 20-fold in Tau^E14^ males. Under induced conditions, Tau^E14^ flies showed lower Tau5A6 signal than Tau^WT^ flies in both females and males. Thus, the stronger behavioural and neurodegenerative effects of Tau^E14^ are not explained by higher total tau abundance and are consistent with enhanced toxicity of the phospho-mimetic variant.

To determine whether early behavioural disruption was accompanied by gross synaptic protein loss, we quantified levels of the postsynaptic scaffolding protein DLG1 and the presynaptic vesicle protein synaptogyrin in whole-head lysates from male and female flies after two weeks of adult-onset tau expression (**Figure 4C,D; Supplementary Figure 6**). DLG1 and synaptogyrin abundance were analysed using Tukey-transformed linear mixed-effects models with Genotype, RU486 status, Sex, and their interactions as fixed effects, and gel number as a random intercept.

For DLG1, there was no significant effect of RU486 status and no significant interactions involving RU486 status or Sex, indicating that tau induction did not detectably alter DLG1 abundance. Although a small main effect of Genotype was detected (χ^2^(2) = 6.28, p = 0.043), planned FDR-adjusted contrasts did not identify induction-dependent changes within tau-expressing genotypes. For synaptogyrin, there were significant main effects of RU486 status (χ^2^(1) = 18.97, p < 0.0001) and Sex (χ^2^(1) = 4.12, p = 0.043), but no significant Genotype × RU486 status interaction or Genotype × RU486 status × Sex interaction. Planned contrasts showed a modest RU-associated increase in synaptogyrin in Tau^WT^ males, but not in Tau^E14^ flies. Thus, early tau-induced behavioural changes were not accompanied by detectable loss of DLG1 or synaptogyrin, suggesting that functional or circuit-level perturbations precede overt depletion of these synaptic markers.

### DAM-based daytime inactivity captures genetic modulation of tau toxicity

Having established a robust female daytime inactivity phenotype following two weeks of adult-onset Tau^E14^ expression, we asked whether this measure could serve as a scalable readout for genetic modulation. We analysed the screen using a Tukey-transformed linear mixed-effects model with Genotype as a fixed effect and Experiment as a random intercept. Candidate modifier conditions were compared with the induced *ElavGS>Tau^E^*^14^ control condition using FDR-adjusted treatment-versus-reference contrasts. A beta mixed model was retained as a sensitivity analysis because the raw sleep/inactivity response is bounded between 0 and 1. **Table 1** shows the candidate genes used in the screen. The primary model showed a strong effect of genotype on daytime inactivity (χ^2^(31) = 978.49, p < 0.0001).

Daytime inactivity showed substantial range across conditions, with clear shifts relative to Tau^E14^ RU+ controls, indicating that this DAM-derived measure is sufficiently sensitive to capture both suppression and enhancement of tau-associated behavioural disruption (Figure 5).

**Figure 5.**
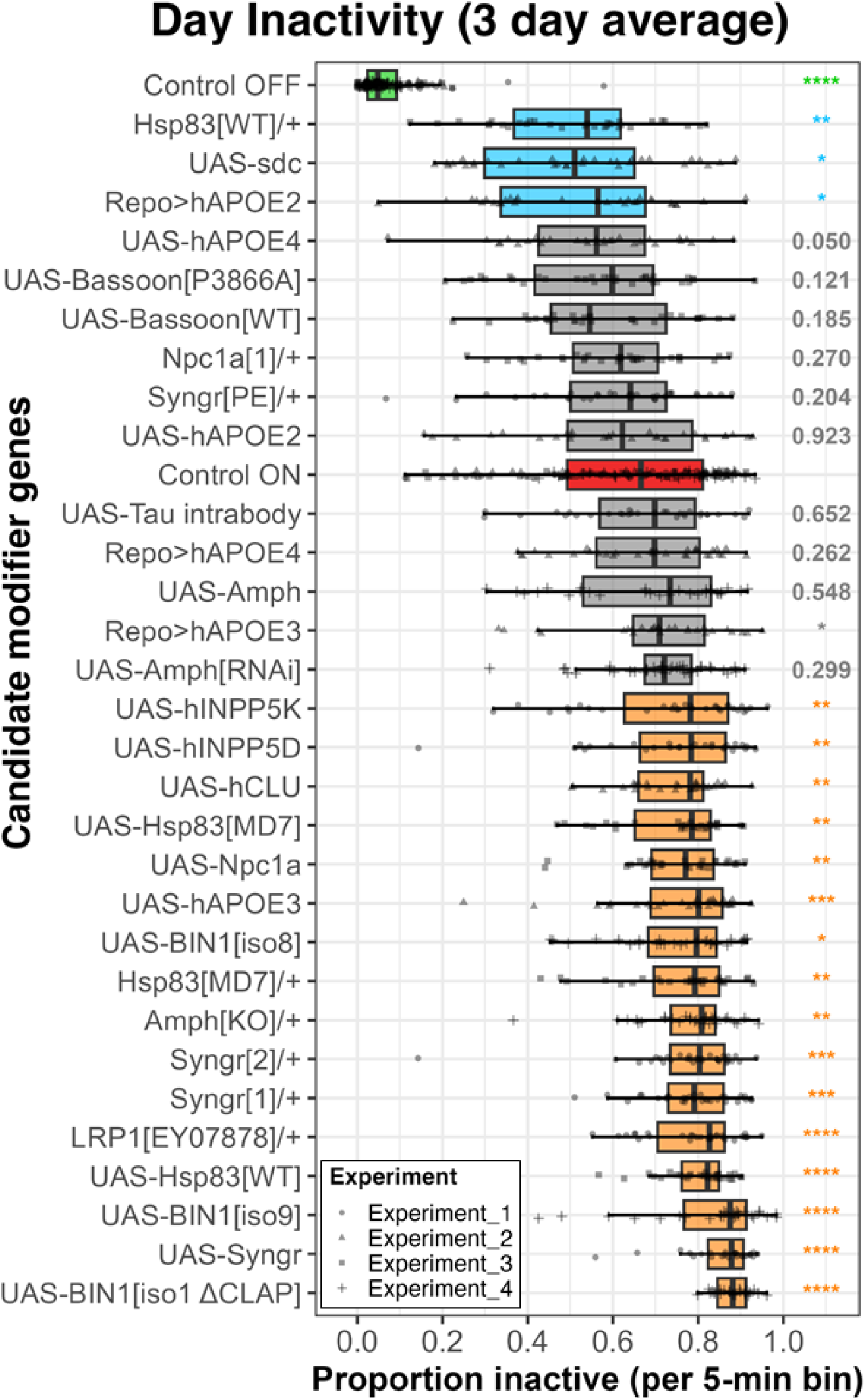
Daytime inactivity across candidate modifier genotypes in a conditional Tau expression screen. Boxplots show the average proportion of time inactive during the daytime over a 3-day recording period for female *ElavGS>Tau^E^*^14^ *Drosophila* expressing candidate modifier genes. The tau-induced control condition (Control ON, RU+) is shown in red and serves as the reference condition for statistical comparisons. A non-induced control (Control OFF, RU-) is shown in green and was included in each experiment to confirm effective induction of Tau expression. All modifier genotypes were analysed under induced conditions (RU+). Modifier genotypes are coloured according to their effect on Tau-induced inactivity: significant suppressors (reduced inactivity relative to Control ON) are shown in blue, significant enhancers (increased inactivity) in orange, and non-significant modifiers in light grey. Individual points represent single flies, with point shapes indicating independent experiments (Experiments 1–4). Genotypes are ordered by mean daytime inactivity. P-values from pairwise comparisons relative to the induced control are displayed to the right of each genotype. Statistical comparisons were performed using a Tukey-transformed linear mixed-effects model with Genotype as a fixed effect and Experiment as a random intercept. Each candidate modifier condition was compared with the induced *ElavGS>Tau^E^*^14^ control condition using FDR-adjusted treatment-versus-reference contrasts. P-values shown are from the primary Tukey-LMM. Modifier genotypes contained *n* = 23–42 flies per condition; control groups comprised *n* = 128 (Control OFF) and *n* = 124 (Control ON). Significance levels are indicated as follows: *P* < 0.05 (\**), P < 0.01 (**), P < 0.001 (\*\*\**), and *P* < 0.0001 (****).

#### Suppressors

Three perturbations significantly suppressed the Tau^E14^-induced daytime inactivity phenotype in the primary model. Glial expression of human APOE2 (Repo>hAPOE2) reduced daytime inactivity relative to induced Tau^E14^ controls, consistent with a protective effect. Neuronal overexpression of syndecan (UAS-sdc) also suppressed the phenotype, in line with evidence implicating heparan sulfate proteoglycans in tau binding, uptake, and toxicity. A third suppressive effect was observed with the genomic Hsp90^WT^ orthologue. This construct reduced daytime inactivity in the primary model, although this effect was weaker in the beta-model sensitivity analysis, suggesting it should be interpreted cautiously. In contrast, other Hsp90 manipulations enhanced the phenotype, indicating context- and construct-dependent effects of this chaperone pathway. This suggests that maintenance of appropriate chaperone balance, rather than simple increases in Hsp90 abundance, may be required to mitigate tau-induced behavioural dysfunction.

#### Enhancers

In contrast, seventeen candidate perturbations significantly enhanced TauE14-induced daytime inactivity in the primary model. These included both overexpression of synaptogyrin (UAS-Syngr) and Syngr loss-of-function alleles, indicating that disruption of synaptogyrin in either direction exacerbates the phenotype and suggesting a requirement for tight regulation of synaptogyrin-dependent synaptic function. BIN1/Amph pathway manipulations also modified the phenotype: human BIN1 isoform 9 and the ΔCLAP isoform strongly enhanced inactivity, with a weaker effect observed for BIN1 isoform 8, while Amph^KO^/+ also enhanced the phenotype. In contrast, UAS-Amph and Amph^RNAi^ did not significantly modify the phenotype under these conditions. Additional enhancers included neuronal expression of human APOE3, glial expression of APOE3, CLU, INPP5D, and the related INPP5K control transgene. Because INPP5K also enhanced the phenotype, the INPP5D result should be interpreted cautiously with respect to gene specificity. Perturbations affecting lipid handling and endocytic pathways also modified the phenotype, including LRP1 loss-of-function and NPC1 overexpression. Within the proteostasis-related group, Hsp90^MD7^/+, UAS-Hsp90^MD7^, and UAS-Hsp90^WT^ enhanced Tau^E14^-induced inactivity, in contrast to the suppressive effect of the genomic Hsp90^WT^ construct. Together with the contrasting effect of Hsp90^WT^, these results suggest a strong dosage- and context-dependence of Hsp90 function in modulating tau toxicity, likely reflecting the requirement for coordinated co-chaperone interactions rather than simple increases in chaperone abundance.

Several additional lines showed no significant effect, including Bassoon variants, UAS-Amph, Amph^RNAi^, neuronal APOE2, Repo>hAPOE4, and Npc1a^1^/+ under the conditions tested. UAS-hAPOE4 did not pass the FDR-corrected significance threshold. These findings indicate that the observed modulation is not simply a general consequence of transgene expression or genetic background.

Together, these results demonstrate that tau-induced behavioural dysfunction can be both suppressed and exacerbated by genetic perturbations, and identify membrane trafficking, lipid metabolism, and proteostasis as key pathways modulating early tau toxicity.

## Discussion

In this study we established that automated behavioural monitoring using the *Drosophila* Activity Monitor (DAM) provides a sensitive readout of tau-driven neurotoxicity in an adult-onset *Drosophila* model. By inducing pan-neuronal expression of human 0N4R tau isoforms specifically in adulthood using GeneSwitch, we showed that both Tau^WT^ and the phospho-mimetic Tau^E14^ are sufficient to shorten lifespan and drive neurodegeneration, while also producing quantifiable disruptions in locomotor activity and inactivity that emerge after induction. These findings reinforce the utility of adult-restricted tau expression for isolating tau-driven pathology from developmental toxicity observed in constitutive models^6,7,13,55^ and position DAM-derived metrics as functionally meaningful outputs that can be measured in a high-throughput manner.

Consistent with the enhanced toxicity of phosphorylation-mimicking tau, Tau^E14^ produced stronger neurodegenerative phenotypes than Tau^WT^, including a greater reduction in lifespan, in agreement with prior studies indicating the potent pathogenicity of hyperphosphorylated tau species^7,56^. While both tau variants induced robust neurodegeneration, differences in vacuolar burden between Tau^WT^ and Tau^E14^ were comparatively modest, suggesting that phosphorylation-mimetic tau may disproportionately impact organismal viability relative to gross structural degeneration at this time point. Vacuolar degeneration was evident across multiple brain regions but was most consistently quantifiable in the optic lobes, providing histopathological evidence of neurodegeneration. This regional vulnerability resembles prior observations in fly tauopathy models^57–59^. Although we detected a significant main effect of sex on vacuole burden, with males generally showing lower vacuole counts than females, we did not detect significant Genotype × Sex, RU486 status × Sex, or Genotype × RU486 status × Sex interactions. Thus, sex influenced overall vacuole burden, but did not strongly modify the induction-dependent tau-associated increase in structural neurodegeneration. These findings suggest that sex can influence the extent of tau-induced neurodegeneration in specific contexts, even in the absence of a strong global effect. Additionally, we observed apparent increases in phalloidin-labelled F-actin in some tau-expressing brains, consistent with the ability of tau toxicity to drive abnormal F-actin stabilisation^60^. Future work will be required to quantify these cytoskeletal changes and determine their relationship to early behavioural disruption.

Behavioural consequences of tau expression were strongly sexually dimorphic. Across assays, females exhibited reduced activity and increased daytime inactivity, whereas males initially became hyperactive and exhibited reduced daytime inactivity. Such divergence suggests that the behavioural consequences of tau expression may reflect interactions between tau-driven neuronal stress and intrinsic biological differences in sleep regulation and arousal circuitry^61–68^. Importantly, this early dimorphism was consistent across RU486 doses and was evident even at the lowest induction levels. Longer-term monitoring indicated that males eventually transition to hypoactivity as neurodegeneration progresses, demonstrating that early hyperactivity is not sustained throughout the lifespan. Notably, although lifespan analyses revealed increased susceptibility of females to tau-induced mortality, vacuolar pathology showed a main effect of sex but no significant sex-by-tau-induction interaction. This indicates that the pronounced early behavioural divergence between sexes is unlikely to be driven solely by differences in gross neurodegeneration, and instead reflects sex-specific responses to tau-induced neuronal dysfunction. This pattern is consistent with evidence for heightened susceptibility to tau pathology in females in vertebrate models^69–71^ and may relate to sex-linked immune and stress-response differences described in *Drosophila*, including stronger NF-κB-driven immune responses regulated by Sex-lethal (Sxl) that can exacerbate neuronal damage^72,73^. Transcriptomic work in tau-null flies further suggests the existence of persistent sex-dependent expression programmes after neural injury^74^, providing a plausible context in which tau-driven stress could yield divergent behavioural and pathological trajectories.

These findings also align with observations in human tauopathies. In Alzheimer’s disease, women show a higher lifetime risk^75^ and, in some studies, greater tau burden and faster cognitive decline at equivalent stages of pathology^76^. Sex differences have also been reported in other tauopathies, including progressive supranuclear palsy^77^ and corticobasal degeneration, although the direction and magnitude of these effects can vary^78^. Proposed mechanisms include the loss of oestrogen-mediated neuroprotection^79^, sex-specific immune responses^80^, and interactions between tau pathology and genetic risk factors such as APOE^81^. The early, behaviourally defined dimorphism observed here may therefore reflect conserved sex-dependent vulnerabilities to tau-induced neuronal dysfunction, supporting the relevance of this model for dissecting mechanisms underlying sex bias in neurodegenerative disease.

Building on the robust female daytime inactivity phenotype, we performed a targeted, proof-of-principle genetic screen using candidate modifiers with established links to Alzheimer’s disease risk and neuronal homeostasis, including BIN1, APOE isoforms, and regulators of membrane trafficking and proteostasis. This revealed substantial bidirectional modulation of tau-induced behavioural dysfunction, demonstrating that this DAM-derived phenotype is both sensitive and biologically informative.

Among suppressors, glial expression of APOE2 reduced Tau^E14^-induced inactivity, consistent with a protective role of this isoform. One plausible explanation is that APOE2 alters lipid transport or extracellular lipidation states in a manner that limits neuronal stress or reduces the propagation of toxic tau species. Given the growing evidence that glial cells regulate neuronal vulnerability through lipid metabolism and extracellular proteostasis, this result supports a model in which non-neuronal mechanisms can modulate early tau-induced dysfunction^82^. Similarly, syndecan overexpression suppressed the phenotype, which may reflect altered heparan sulfate–mediated interactions at the neuronal surface. As heparan sulfate proteoglycans have been implicated in tau binding, uptake, and spread, increased syndecan levels could modify tau internalisation dynamics or sequester extracellular tau, thereby reducing its toxic impact^83^.

In contrast, a larger set of modifiers enhanced tau-induced inactivity. Notably, both overexpression and loss-of-function of synaptogyrin exacerbated the phenotype, suggesting that disruption of synaptic vesicle organisation in either direction sensitises neurons to tau toxicity^10,40^. This bidirectional effect argues against a simple gain- or loss-of-function mechanism and instead points to a requirement for tight homeostatic control of synaptogyrin levels. Similarly, multiple BIN1 isoforms enhanced the phenotype, consistent with proposed roles for BIN1 in membrane curvature, endocytosis, and tau interaction, and suggesting that altered membrane dynamics may potentiate tau-induced dysfunction^45^.

Additional enhancers included neuronal and glial APOE3, CLU, INPP5D, and the related INPP5K control transgene, as well as perturbations affecting lipid handling and endocytic pathways such as LRP1 loss-of-function and NPC1 overexpression. The parallel effect of INPP5K suggests that the INPP5D result should be interpreted cautiously until validated with additional reagents. These collectively point toward membrane trafficking and lipid metabolism as key modulators of early tau toxicity, potentially by influencing tau localisation, clearance, or access to vulnerable neuronal compartments. Perturbation of proteostasis pathways also had complex effects: while near-physiological expression of Hsp90^WT^ was protective, multiple other Hsp90-related constructs enhanced the phenotype. This divergence suggests that tau toxicity is sensitive to the balance of chaperone systems, where disruption of co-chaperone interactions or stoichiometry may be more detrimental than modest restoration of baseline Hsp90 levels^84^.

An important limitation of this study is that modifier effects were assessed only in the presence of Tau^E14^. As such, it remains possible that some “enhancers” reflect general declines in organismal health, locomotor capacity, or neuronal function independent of tau. Future experiments comparing these lines in the absence of tau expression will be necessary to distinguish tau-specific modifiers from those that broadly impair behavioural output.

Collectively, these findings suggest that early tau-induced behavioural dysfunction is highly sensitive to perturbations in lipid metabolism, membrane trafficking, synaptic homeostasis, and proteostasis. More broadly, they demonstrate that DAM-based behavioural assays provide a scalable platform for identifying and prioritising candidate modifiers in vivo, while also highlighting the need for careful secondary validation to resolve tau-specific mechanisms.

Several other limitations should be noted. RU486 itself can influence physiology and behaviour, including lifespan and gut biology in females^85^, although effects on activity and sleep are not consistently observed^86^. In our experiments, tau-dependent phenotypes were distinct from, and often exceeded, RU486-only effects, but careful use of matched induced and uninduced controls remains important. Expression of human tau via the UAS system may also result in supraphysiological levels relative to endogenous conditions, and vacuolar pathology represents a relatively coarse measure of neurodegeneration compared to stereological approaches. More broadly, not all aspects of human tau pathology are fully recapitulated in *Drosophila;* for example, evidence for trans-synaptic tau propagation remains mixed, with studies reporting both the presence and absence of neuronal spread^34,87,88^. Finally, the targeted genetic screen was performed in females to maximise sensitivity to a robust behavioural phenotype, and future work will be needed to assess the generality of these modifiers across sexes^89^. Despite these considerations, the consistency of behavioural, lifespan, and neurodegenerative phenotypes supports the utility of this system, which provides a tractable platform for dissecting early tau toxicity and its modulation across biological contexts.

In conclusion, this study provides a model of adult-onset tau toxicity in *Drosophila* and demonstrates that DAM-based behavioural analysis captures quantifiable signatures of neuronal dysfunction. Combining temporally controlled tau induction with organismal, histological, and behavioural outcomes enables detection of early functional deficits and supports the use of DAM phenotyping for high-throughput screening of tau phenotypes.

Future studies extending screens, integrating more precise behavioural and physiological measurements, and validating conserved modifiers in human tissue will be essential for using this model to understand and ultimately intervene in tauopathy progression.

## Acknowledgements

We are grateful to past and present members of the Durrant and Spires-Jones labs for helpful discussions. We thank Dr Adam Dobson (University of Glasgow) for the CoxPH lifespan analysis R script. We also thank Michael Molinek and Krisztina Vinko for management and delivery of the *Drosophila* food and Jane Tulloch for administrative support. We also thank the wider fly community for the generous sharing of reagents and stocks, particularly the Bloomington *Drosophila* Stock Center (NIH P40OD018537). Cartoon in Figure 1 was generated using BioRender.

## Data availability

All relevant data can be found within the article and its supplementary information and will be available in Edinburgh University Data Repository.

## Supplementary data

**Supplementary Figure 1.**
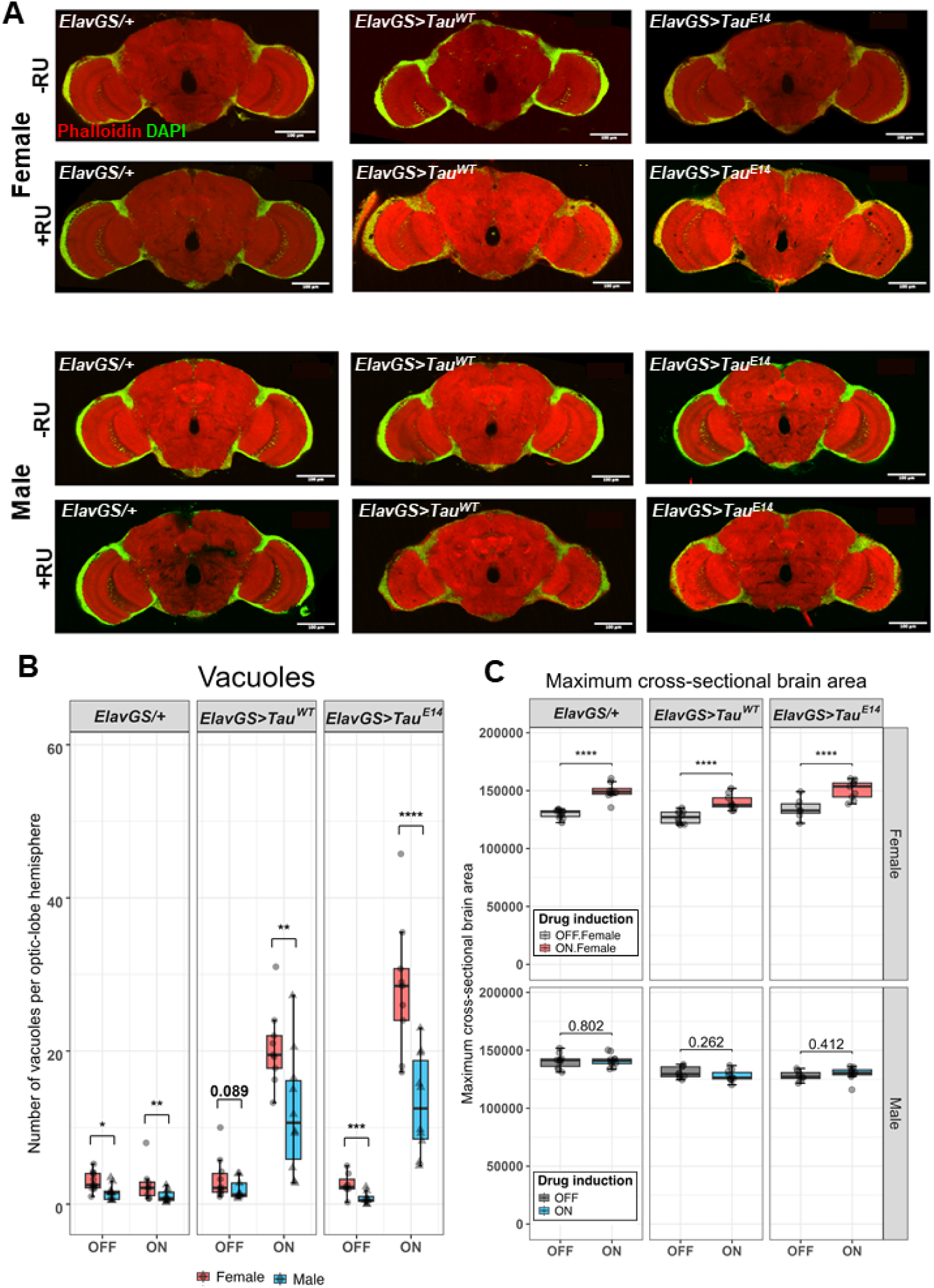
Whole-brain imaging, sex-stratified vacuole analysis, and brain-area quantification after four weeks of adult-onset tau expression. (**A**) Representative 2-photon micrographs of whole-mount adult fly brains corresponding to Figure 1D. Top panels show females and bottom panels show males, with rows indicating induction status (−RU or +RU). Brains were stained with phalloidin to visualise neuropil structure and DAPI to label cell nuclei. Scale bar: 100 µm. (**B**) Sex-stratified quantification of vacuole counts within each Genotype × RU486 status condition. Points show individual brains/images, with counts averaged across scorers for visualisation. Statistical comparisons show Male versus Female contrasts within each Genotype × RU486 status stratum from the primary negative binomial mixed-effects model. P-values were adjusted using the false discovery rate method. (**C**) Quantification of maximum cross-sectional brain area from 2-photon micrographs. Points show individual brains/images, with measurements averaged across scorers for visualisation. Brain area was analysed using a Gamma mixed-effects model with Genotype, RU486 status, Sex, and their interactions as fixed effects, scorer as a fixed effect, and ImageID as a random intercept. RU486 induction significantly increased brain area in females across genotypes but not in males. Significance thresholds are indicated as P < 0.05 (*), P < 0.01 (**), P < 0.001 (***), and P < 0.0001 (****).

**Supplementary Figure 2.**
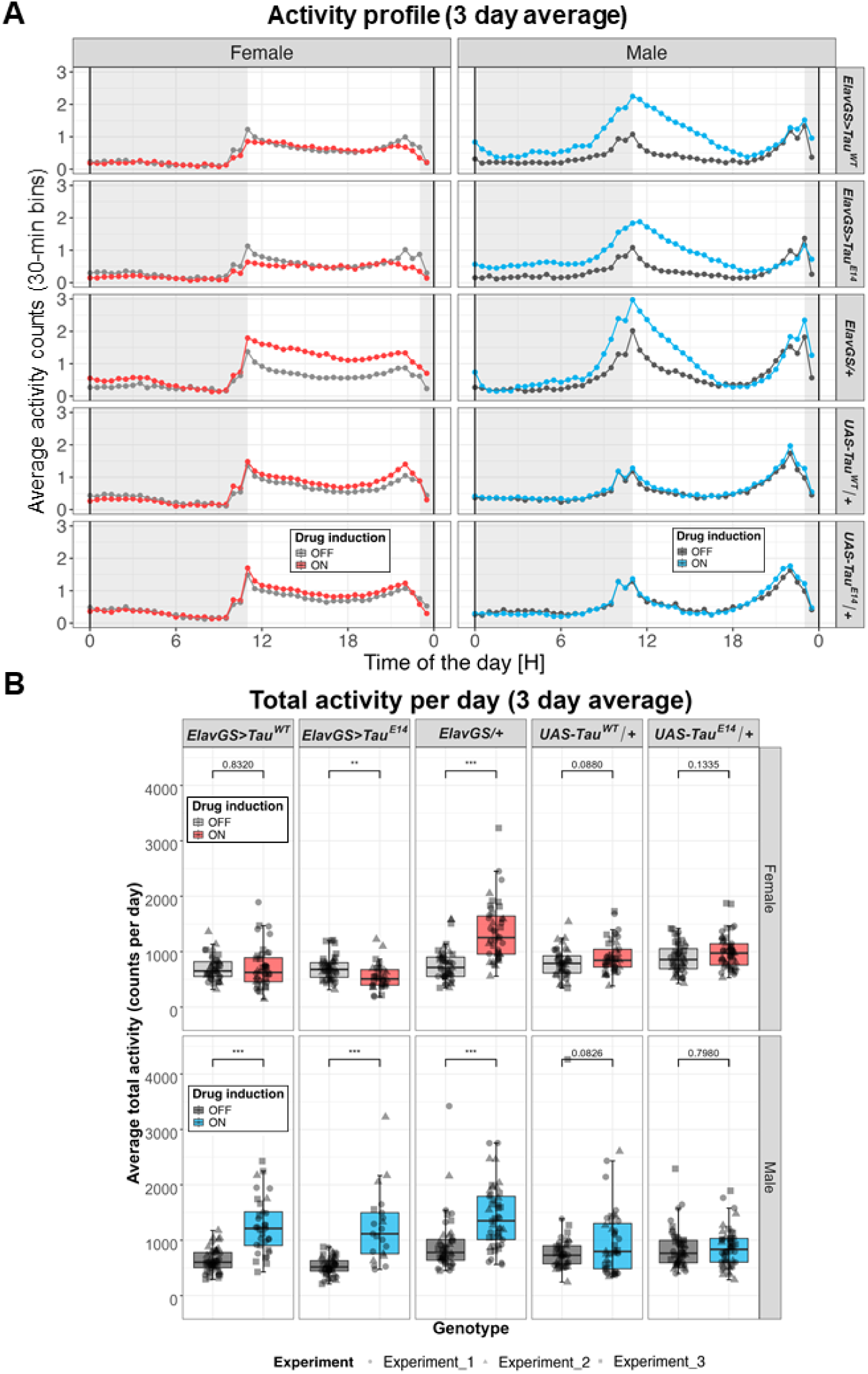
Adult-onset pan-neuronal expression of human Tau for 2 weeks alters locomotor activity in a sexually dimorphic manner. (**A**) Average daily locomotor activity profiles (3-day mean) under 12:12 light–dark conditions showing population mean activity counts across the day. Activity counts were averaged in 30-minute bins for each genotype. Shaded regions indicate the dark phase. (**B**) Boxplots showing average daily locomotor activity under LD conditions. Adult-onset pan-neuronal expression of Tau^WT^ or Tau^E14^ resulted in increased locomotor activity in males and reduced activity in females relative to controls. Individual points represent single flies and point shapes indicate independent experiments. Boxplot colours denote RU486 drug status: grey (–RU; no drug, Tau expression OFF) and coloured (+RU; drug administered, Tau expression ON; red = females, blue = males). Flies were 2-weeks old when introduced into the DAMS. Statistical analysis of daily locomotor activity was performed using a negative binomial mixed-effects model with Genotype, RU486 status, Sex, and their interactions as fixed effects, and Experiment as a random intercept. Pairwise comparisons show planned +RU versus −RU contrasts within each Genotype × Sex stratum. P-values are derived from the negative binomial model. Significance thresholds are indicated as *P* < 0.05 (\**), P < 0.01 (**), and P < 0.001 (\*\*\**). Sample sizes ranged from *n* = 21–48 flies per condition.

**Supplementary Figure 3.**
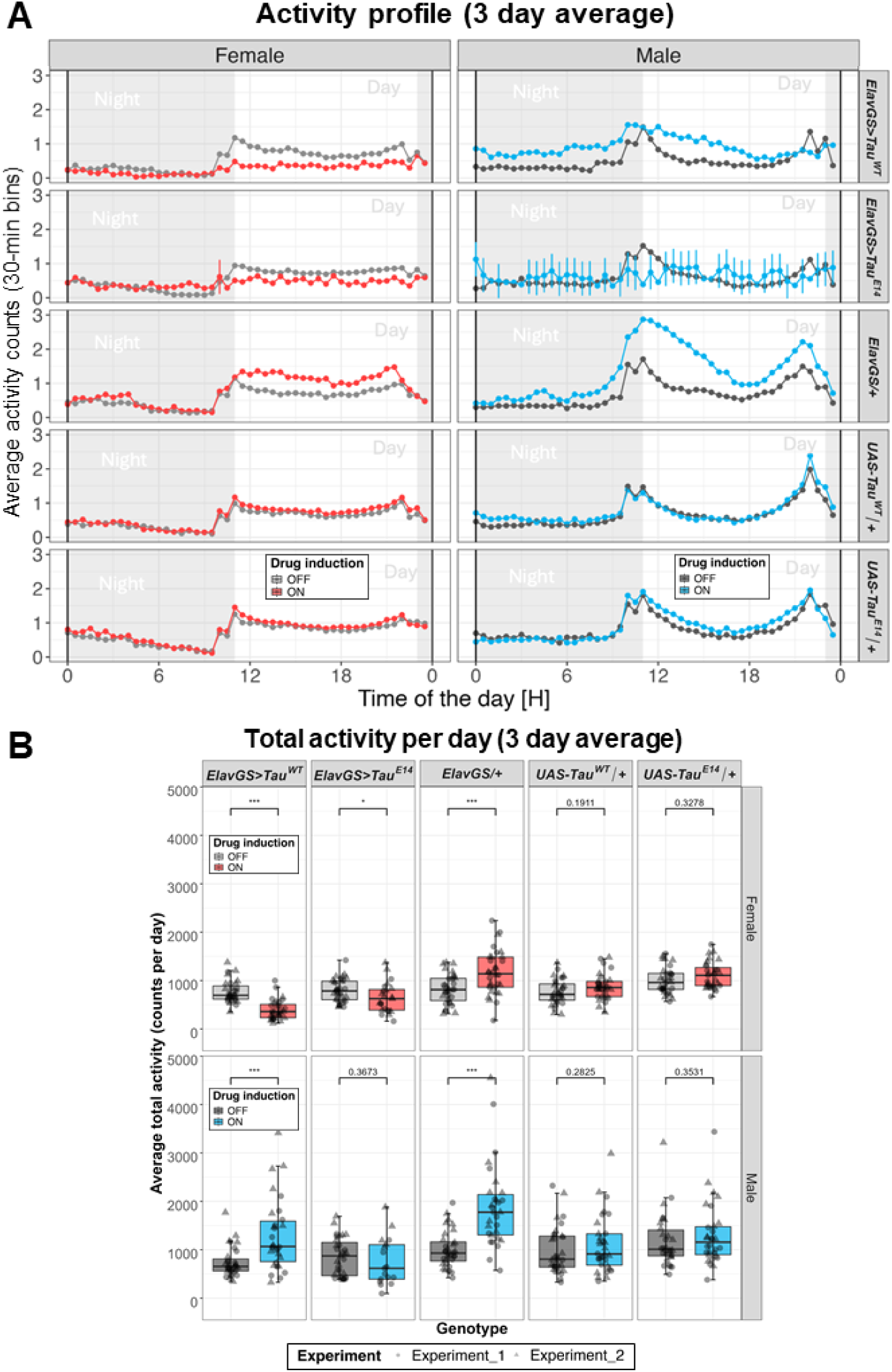
Prolonged adult-onset Tau expression alters locomotor activity after four weeks. (**A**) Average daily locomotor activity profiles under 12:12 light–dark conditions showing population mean activity counts across the day. Activity counts were averaged in 30-minute bins. Profiles represent data averaged from two independent experiments. Flies were 4 weeks old when introduced into the DAMS. Shaded regions indicate the dark phase. (**B**) Boxplots showing average daily locomotor activity under LD conditions. Individual points represent single flies and point shapes indicate independent experiments. Boxplot colours denote RU486 drug status: grey (−RU; no drug administered, Tau expression OFF) and coloured (+RU; drug administered, Tau expression ON; red = females, blue = males). Statistical analysis was performed using a negative binomial mixed-effects model with Genotype, RU486 status, Sex, and their interactions as fixed effects, and Experiment as a random intercept. Pairwise comparisons show planned +RU versus −RU contrasts within each Genotype × Sex stratum. P-values are derived from the negative binomial model. After four weeks, RU486 induction reduced activity in *ElavGS>Tau^WT^* and *ElavGS>Tau^E^*^14^ females, increased activity in *ElavGS>Tau^WT^*males, and did not significantly alter total daily activity in *ElavGS>Tau^E^*^14^ males. RU486 also increased activity in *ElavGS/+* controls. Significance thresholds are indicated as P < 0.05 (*), P < 0.01 (**), and P < 0.001 (***). Sample sizes ranged from n = 17–32 flies per condition.

**Supplementary Figure 4.**
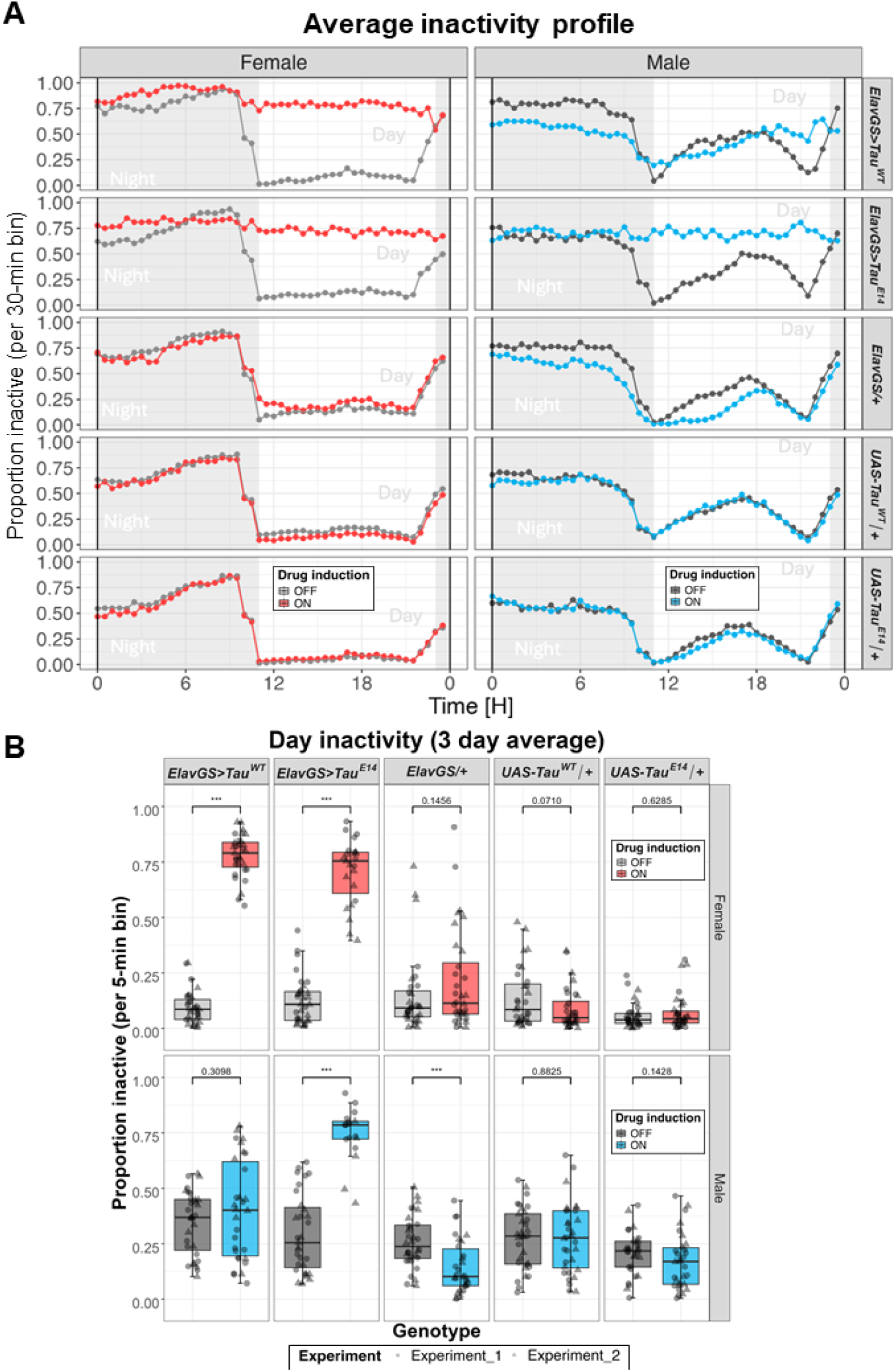
Prolonged adult-onset Tau expression alters daytime sleep/inactivity after four weeks. (**A**) Average sleep/inactivity profiles under 12:12 light–dark conditions showing population mean inactivity across the day. Sleep/inactivity values were averaged in 30-minute bins. Profiles represent data averaged from two independent experiments. Flies were 4 weeks old when introduced into the DAMS. Shaded regions indicate the dark phase. (**B**) Boxplots showing mean daytime sleep/inactivity across genotypes and RU486 conditions. Individual points represent single flies and point shapes indicate independent experiments. Boxplot colours denote RU486 drug status: grey (−RU; no drug administered, Tau expression OFF) and coloured (+RU; drug administered, Tau expression ON; red = females, blue = males). Statistical analysis was performed using a beta mixed-effects model with Genotype, RU486 status, Sex, and their interactions as fixed effects, and Experiment as a random intercept. Pairwise comparisons show planned +RU versus −RU contrasts within each Genotype × Sex stratum. P-values are derived from the beta model. After four weeks, RU486 induction increased daytime sleep/inactivity in *ElavGS>Tau^WT^* and *ElavGS>Tau^E^*^14^ females and in *ElavGS>Tau^E^*^14^ males, but not in *ElavGS>Tau^WT^* males. RU486 reduced daytime sleep/inactivity in male *ElavGS/+* controls. Significance thresholds are indicated as P < 0.05 (*), P < 0.01 (**), and P < 0.001 (***). Sample sizes ranged from n = 17–32 flies per condition.

**Supplementary Figure 5.**
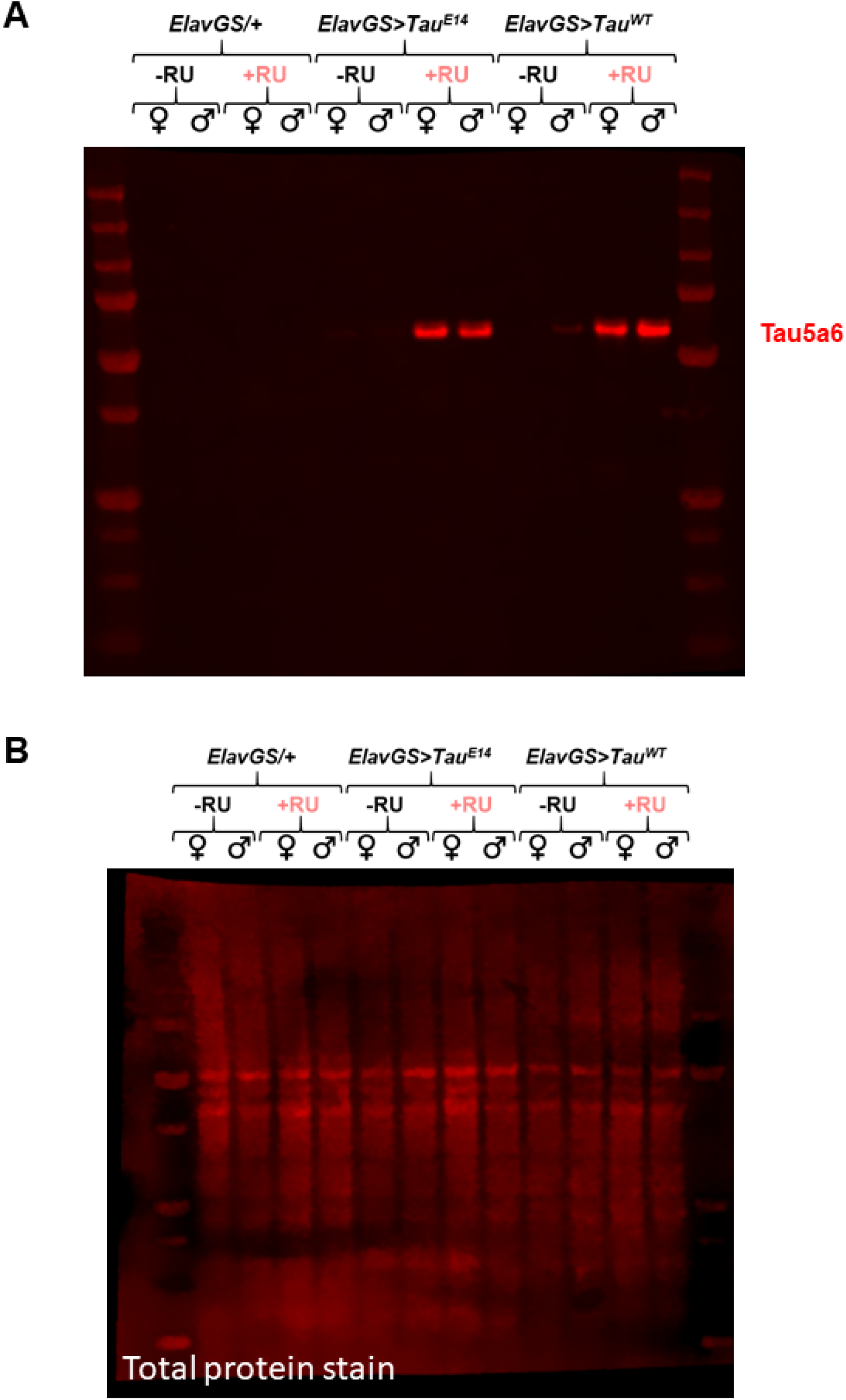
Full western blots for Tau protein analysis. (**A**) Full-length western blots corresponding to Figure 4A showing Tau5A6 signal in whole-head lysates after two weeks of adult-onset tau expression. Samples include *ElavGS/+, ElavGS>Tau^WT^*, and *ElavGS>Tau^E^*^14^ flies under induced (+RU) and uninduced (−RU) conditions in both sexes. (**B**) Corresponding total protein stain used for normalisation. Total protein signal was used to control for loading variability across samples.

**Supplementary Figure 6.**
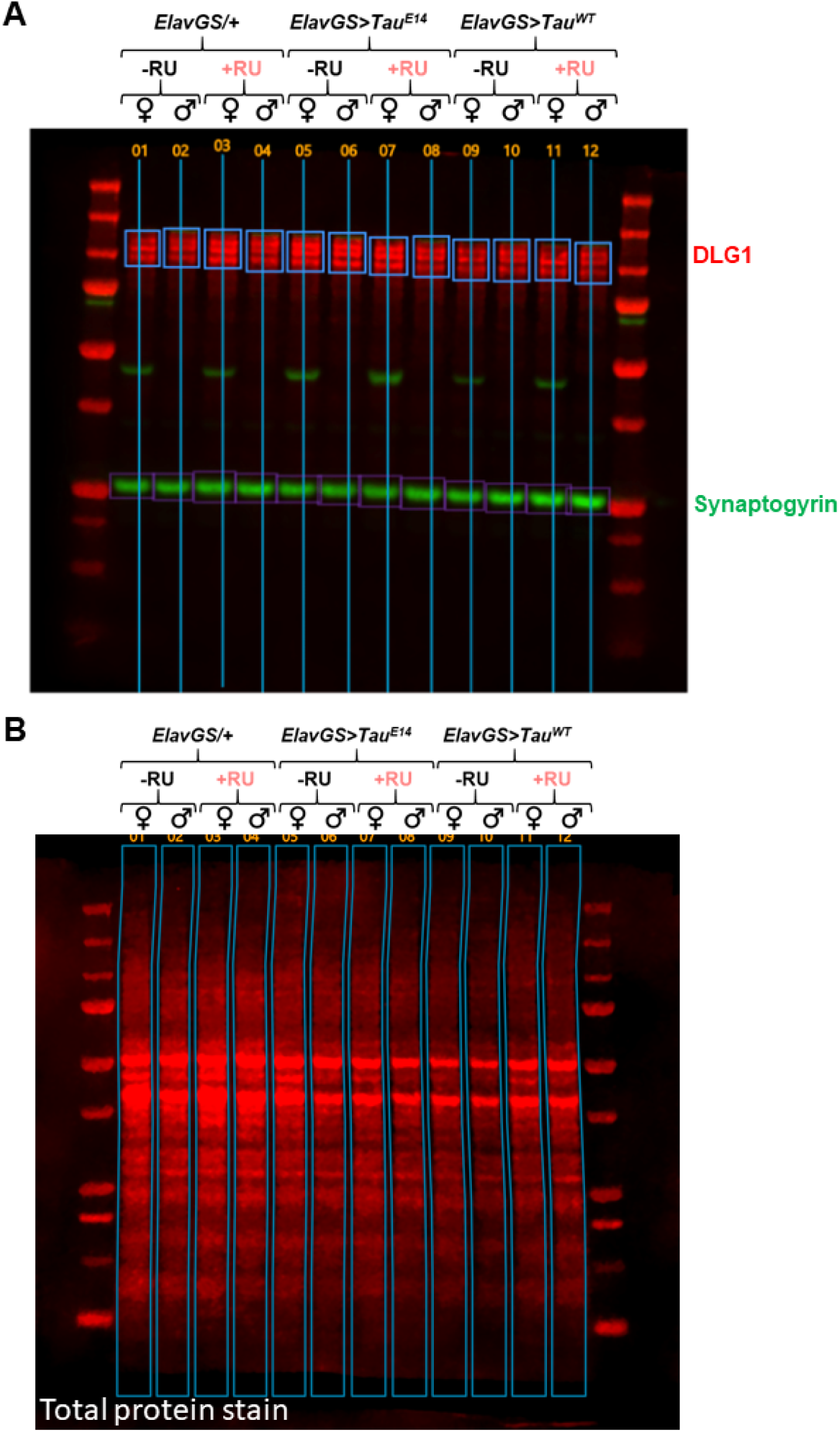
Full western blots for synaptic protein analysis. (**A**) Representative full-length western blots corresponding to Figure 4C showing DLG1 (red; higher molecular weight) and synaptogyrin (green; lower molecular weight) in whole-head lysates from flies following two weeks of adult-onset tau expression. Samples include Tau^WT^ and Tau^E14^ under induced (+RU) and uninduced (−RU) conditions, alongside appropriate controls. (**B**) Corresponding total protein stain for all lanes used for normalisation. Total protein signal was used to control for loading variability across samples.

## References

1. Kovacs GG. Chapter 25 - Tauopathies. In: Kovacs GG, Alafuzoff I, eds. Handbook of Clinical Neurology. Vol 145. Neuropathology. Elsevier; 2018:355–368. doi:10.1016/B978-0-12-802395-2.00025-0

2. Lane-Donovan C, Boxer AL. Disentangling tau: One protein, many therapeutic approaches. Neurotherapeutics. 2024;21(2):e00321. doi:10.1016/j.neurot.2024.e00321

3. Braak H, Thal DR, Ghebremedhin E, Del Tredici K. Stages of the pathologic process in Alzheimer disease: age categories from 1 to 100 years. J Neuropathol Exp Neurol. 2011;70:960–969.

4. Wang Y, Mandelkow E. Tau in physiology and pathology. Nat Rev Neurosci. 2016;17(1):22–35. doi:10.1038/nrn.2015.1

5. Sivanantharajah L, Mudher A, Shepherd D. An evaluation of Drosophila as a model system for studying tauopathies such as Alzheimer’s disease. Journal of Neuroscience Methods. 2019;319:77–88. doi:10.1016/j.jneumeth.2019.01.001

6. Wittmann CW, Wszolek MF, Shulman JM, et al. Tauopathy in Drosophila: neurodegeneration without neurofibrillary tangles. Science. 2001;293:711–714.

7. Khurana V, Lu Y, Steinhilb ML, Oldham S, Shulman JM, Feany MB. TOR-Mediated Cell-Cycle Activation Causes Neurodegeneration in a Drosophila Tauopathy Model. Current Biology. 2006;16(3):230–241. doi:10.1016/j.cub.2005.12.042

8. Moloney A, Sattelle DB, Lomas DA, Crowther DC. Alzheimer’s disease: insights from Drosophila melanogaster models. Trends Biochem Sci. 2010;35(4):228–235. doi:10.1016/j.tibs.2009.11.004

9. Keramidis I, Vourkou E, Papanikolopoulou K, Skoulakis EMC. Functional Interactions of Tau Phosphorylation Sites That Mediate Toxicity and Deficient Learning in Drosophila melanogaster. Front Mol Neurosci. 2020;13. doi:10.3389/fnmol.2020.569520

10. Largo-Barrientos P, Apóstolo N, Creemers E, et al. Lowering Synaptogyrin-3 expression rescues Tau-induced memory defects and synaptic loss in the presence of microglial activation. Neuron. 2021;109(5):767–777.e5. doi:10.1016/j.neuron.2020.12.016

11. Pfeiffenberger C, Lear BC, Keegan KP, Allada R. Locomotor Activity Level Monitoring Using the Drosophila Activity Monitoring (DAM) System. Cold Spring Harb Protoc. 2010;2010(11):pdb.prot5518. doi:10.1101/pdb.prot5518

12. Cichewicz K, Hirsh J. ShinyR-DAM: a program analyzing Drosophila activity, sleep and circadian rhythms. Commun Biol. 2018;1(1):1–5. doi:10.1038/s42003-018-0031-9

13. Buhl E, Higham JP, Hodge JJL. Alzheimer’s disease-associated tau alters Drosophila circadian activity, sleep and clock neuron electrophysiology. Neurobiology of Disease. 2019;130:104507. doi:10.1016/j.nbd.2019.104507

14. Woods JK, Kowalski S, Rogina B. Determination of the Spontaneous Locomotor Activity in Drosophila melanogaster. J Vis Exp. 2014;(86):51449. doi:10.3791/51449

15. Mershin A, Pavlopoulos E, Fitch O, Braden BC, Nanopoulos DV, Skoulakis EMC. Learning and Memory Deficits Upon TAU Accumulation in Drosophila Mushroom Body Neurons. Learn Mem. 2004;11(3):277–287. doi:10.1101/lm.70804

16. Higham JP, Malik BR, Buhl E, et al. Alzheimer’s Disease Associated Genes Ankyrin and Tau Cause Shortened Lifespan and Memory Loss in Drosophila. Front Cell Neurosci. 2019;13. doi:10.3389/fncel.2019.00260

17. Gozes I, Shapira G, Lobyntseva A, Shomron N. Unexpected gender differences in progressive supranuclear palsy reveal efficacy for davunetide in women. Transl Psychiatry. 2023;13(1):319. doi:10.1038/s41398-023-02618-9

18. Andrews D, Ducharme S, Chertkow H, Sormani MP, Collins DL, Initiative for the ADN. The higher benefit of lecanemab in males compared to females in CLARITY AD is probably due to a real sex effect. Alzheimer’s & Dementia. 2025;21(1):e14467. doi:10.1002/alz.14467

19. Zeighami Y, Tremblay C, Dadar M. Sex differences in vulnerability to tau pathology: Impact on cognitive decline. Alzheimers Dement. 2025;21(10):e70634. doi:10.1002/alz.70634

20. Smith R, Strandberg O, Mattsson-Carlgren N, et al. The accumulation rate of tau aggregates is higher in females and younger amyloid-positive subjects. Brain. 2020;143(12):3805–3815. doi:10.1093/brain/awaa327

21. Vourkou E, Paspaliaris V, Bourouliti A, et al. Differential Effects of Human Tau Isoforms to Neuronal Dysfunction and Toxicity in the Drosophila CNS. Int J Mol Sci. 2022;23(21):12985. doi:10.3390/ijms232112985

22. Cassar M, Law AD, Chow ES, Giebultowicz JM, Kretzschmar D. Disease-Associated Mutant Tau Prevents Circadian Changes in the Cytoskeleton of Central Pacemaker Neurons. Front Neurosci. 2020;14. doi:10.3389/fnins.2020.00232

23. Jaciuch D, Munns J, Chawla S, Davis SJ, Juusola M. Selective human tau protein expression in different clock circuits of the Drosophila brain disrupts different aspects of sleep and circadian rhythms. bioRxiv. Preprint posted online December 14, 2020:2020.12.14.422675. doi:10.1101/2020.12.14.422675

24. Revel M, Nagoshi E, Maeda R. Drosophila video-assisted activity monitor (DrosoVAM): a versatile method for behaviour monitoring. R Soc Open Sci. 12(9):250764. doi:10.1098/rsos.250764

25. Akpoghiran O, Strich AK, Koh K. Effects of sex, mating status, and genetic background on circadian behavior in Drosophila. Front Neurosci. 2025;18. doi:10.3389/fnins.2024.1532868

26. Woodling N. Sex- and strain-dependent effects of ageing on sleep and activity patterns in Drosophila. PLOS ONE. 2024;19(8):e0308652. doi:10.1371/journal.pone.0308652

27. Goodman LD, Ralhan I, Li X, et al. Tau is required for glial lipid droplet formation and resistance to neuronal oxidative stress. Nat Neurosci. 2024;27(10):1918–1933. doi:10.1038/s41593-024-01740-1

28. Bukhari H, Nithianandam V, Battaglia RA, et al. Transcriptional programs mediating neuronal toxicity and altered glial–neuronal signaling in a Drosophila knock-in tauopathy model. Genome Res. 2024;34(4):590–605. doi:10.1101/gr.278576.123

29. Leventhal MJ, Zanella CA, Kang B, et al. An integrative systems-biology approach defines mechanisms of Alzheimer’s disease neurodegeneration. Nat Commun. 2025;16:4441. doi:10.1038/s41467-025-59654-w

30. Osterwalder T, Yoon KS, White BH, Keshishian H. A conditional tissue-specific transgene expression system using inducible GAL4. Proceedings of the National Academy of Sciences. 2001;98(22):12596–12601. doi:10.1073/pnas.221303298

31. Gorsky MK, Burnouf S, Dols J, Mandelkow E, Partridge L. Acetylation mimic of lysine 280 exacerbates human Tau neurotoxicity in vivo. Sci Rep. 2016;6(1):22685. doi:10.1038/srep22685

32. Catterson JH, Minkley L, Aspe S, et al. Protein retention in the endoplasmic reticulum rescues Aβ toxicity in Drosophila. Neurobiology of Aging. 2023;132:154–174. doi:10.1016/j.neurobiolaging.2023.09.008

33. Latouche M, Lasbleiz C, Martin E, et al. A Conditional Pan-Neuronal Drosophila Model of Spinocerebellar Ataxia 7 with a Reversible Adult Phenotype Suitable for Identifying Modifier Genes. J Neurosci. 2007;27(10):2483–2492. doi:10.1523/JNEUROSCI.5453-06.2007

34. Catterson JH, Mouofo EN, López De Toledo Soler I, et al. Drosophila appear resistant to trans-synaptic tau propagation. Brain Communications. 2024;6(4):fcae256. doi:10.1093/braincomms/fcae256

35. Williams DM, Heikkinen S, Hiltunen M, Davies NM, Anderson EL. The proportion of Alzheimer’s disease attributable to apolipoprotein E. npj Dement. 2026;2(1):1. doi:10.1038/s44400-025-00045-9

36. Marvian AT, Strauss T, Tang Q, et al. Distinct regulation of Tau Monomer and aggregate uptake and intracellular accumulation in human neurons. Mol Neurodegeneration. 2024;19(1):100. doi:10.1186/s13024-024-00786-w

37. Tuck BJ, Miller LVC, Katsinelos T, et al. Cholesterol determines the cytosolic entry and seeded aggregation of tau. Cell Reports. 2022;39(5):110776. doi:10.1016/j.celrep.2022.110776

38. Martinez P, Patel H, You Y, et al. Bassoon contributes to tau-seed propagation and neurotoxicity. Nat Neurosci. 2022;25(12):1597–1607. doi:10.1038/s41593-022-01191-6

39. Stevens RJ, Akbergenova Y, Jorquera RA, Littleton JT. Abnormal Synaptic Vesicle Biogenesis in Drosophila Synaptogyrin Mutants. J Neurosci. 2012;32(50):18054–18067. doi:10.1523/JNEUROSCI.2668-12.2012

40. McInnes J, Wierda K, Snellinx A, et al. Synaptogyrin-3 Mediates Presynaptic Dysfunction Induced by Tau. Neuron. 2018;97(4):823–835.e8. doi:10.1016/j.neuron.2018.01.022

41. Bellenguez C, Küçükali F, Jansen IE, et al. New insights into the genetic etiology of Alzheimer’s disease and related dementias. Nat Genet. 2022;54(4):412–436. doi:10.1038/s41588-022-01024-z

42. Levy SA, Zuniga G, Gonzalez EM, Butler D, Temple S, Frost B. TauLUM, an in vivo Drosophila sensor of tau multimerization, identifies neuroprotective interventions in tauopathy. Cell Rep Methods. 2022;2(9):100292. doi:10.1016/j.crmeth.2022.100292

43. Rauch JN, Luna G, Guzman E, et al. LRP1 is a master regulator of tau uptake and spread. Nature. 2020;580:381–385.

44. Ben Khalaf N. Heat shock proteins (Hsp70 and Hsp90) in neurodegeneration: pathogenic roles and therapeutic potential. Front Aging Neurosci. 18:1711422. doi:10.3389/fnagi.2026.1711422

45. Lauwers E, Wang YC, Gallardo R, et al. Hsp90 Mediates Membrane Deformation and Exosome Release. Molecular Cell. 2018;71(5):689–702.e9. doi:10.1016/j.molcel.2018.07.016

46. Bass TM, Grandison RC, Wong R, Martinez P, Partridge L, Piper MDW. Optimization of Dietary Restriction Protocols in Drosophila. J Gerontol A Biol Sci Med Sci. 2007;62(10):1071–1081. doi:10.1093/gerona/62.10.1071

47. Behnke JA, Ye C, Moberg KH, Zheng JQ. A protocol to detect neurodegeneration in Drosophila melanogaster whole-brain mounts using advanced microscopy. STAR Protocols. 2021;2(3):100689. doi:10.1016/j.xpro.2021.100689

48. Schindelin J, Arganda-Carreras I, Frise E, et al. Fiji: an open-source platform for biological-image analysis. Nat Methods. 2012;9(7):676–682. doi:10.1038/nmeth.2019

49. Hendricks JC, Finn SM, Panckeri KA, et al. Rest in Drosophila Is a Sleep-like State. Neuron. 2000;25(1):129–138. doi:10.1016/S0896-6273(00)80877-6

50. Shaw PJ, Cirelli C, Greenspan RJ, Tononi G. Correlates of Sleep and Waking in Drosophila melanogaster. Science. 2000;287(5459):1834–1837. doi:10.1126/science.287.5459.1834

51. Nitz DA, van Swinderen B, Tononi G, Greenspan RJ. Electrophysiological Correlates of Rest and Activity in *Drosophila melanogaster*. Current Biology. 2002;12(22):1934–1940. doi:10.1016/S0960-9822(02)01300-3

52. Vecsey CG, Koochagian C, Reyes M, Sitaraman D. Neural Stimulation during Drosophila Activity Monitor (DAM)-Based Studies of Sleep and Circadian Rhythms in Drosophila melanogaster. Cold Spring Harb Protoc. 2024;2024(11):pdb.prot108180. doi:10.1101/pdb.prot108180

53. Posit team. RStudio: Integrated Development Environment for R. Published online 2025. http://www.posit.co/.

54. Piper MD. Piper Lab Resources: Excel Lifespan Sheets. PiperLab. 2026. Accessed March 12, 2026. https://piperlab.org/resources/

55. Fernius J, Starkenberg A, Pokrzywa M, Thor S. Human TTBK1, TTBK2 and MARK1 kinase toxicity in Drosophila melanogaster is exacerbated by co-expression of human Tau. Biology Open. 2017;6(7):1013–1023. doi:10.1242/bio.022749

56. Abtahi SL, Masoudi R, Haddadi M. The distinctive role of tau and amyloid beta in mitochondrial dysfunction through alteration in Mfn2 and Drp1 mRNA Levels: A comparative study in Drosophila melanogaster. Gene. 2020;754:144854. doi:10.1016/j.gene.2020.144854

57. Barati A, Masoudi R, Yousefi R, Monsefi M, Mirshafiey A. Tau and amyloid beta differentially affect the innate immune genes expression in Drosophila models of Alzheimer’s disease and β- D Mannuronic acid (M2000) modulates the dysregulation. Gene. 2022;808:145972. doi:10.1016/j.gene.2021.145972

58. Passarella D, Goedert M. Beta-sheet assembly of Tau and neurodegeneration in Drosophila melanogaster. Neurobiology of Aging. 2018;72:98–105. doi:10.1016/j.neurobiolaging.2018.07.022

59. Oliveira AC, Santos M, Pinho M, Lopes CS. String/Cdc25 phosphatase is a suppressor of Tau-associated neurodegeneration. Disease Models & Mechanisms. 2023;16(1):dmm049693. doi:10.1242/dmm.049693

60. Fulga TA, Elson-Schwab I, Khurana V, et al. Abnormal bundling and accumulation of F-actin mediates tau-induced neuronal degeneration in vivo. Nat Cell Biol. 2007;9(2):139–148. doi:10.1038/ncb1528

61. Andretic R, Shaw PJ. Essentials of Sleep Recordings in *Drosophila:* Moving Beyond Sleep Time. In: Young MW, ed. Methods in Enzymology. Vol 393. Circadian Rhythms. Academic Press; 2005:759–772. doi:10.1016/S0076-6879(05)93040-1

62. Isaac RE, Li C, Leedale AE, Shirras AD. Drosophila male sex peptide inhibits siesta sleep and promotes locomotor activity in the post-mated female. Proceedings of the Royal Society B: Biological Sciences. 2009;277(1678):65–70. doi:10.1098/rspb.2009.1236

63. Zhang W, Guo C, Chen D, Peng Q, Pan Y. Hierarchical Control of Drosophila Sleep, Courtship, and Feeding Behaviors by Male-Specific P1 Neurons. Neurosci Bull. 2018;34(6):1105–1110. doi:10.1007/s12264-018-0281-z

64. Machado DR, Afonso DJ, Kenny AR, et al. Identification of octopaminergic neurons that modulate sleep suppression by male sex drive. Griffith LC, ed. eLife. 2017;6:e23130. doi:10.7554/eLife.23130

65. Toda H, Williams JA, Gulledge M, Sehgal A. A sleep-inducing gene, nemuri, links sleep and immune function in Drosophila. Science. 2019;363(6426):509–515. doi:10.1126/science.aat1650

66. Wu B, Ma L, Zhang E, et al. Sexual dimorphism of sleep regulated by juvenile hormone signaling in Drosophila. PLoS Genet. 2018;14(4):e1007318. doi:10.1371/journal.pgen.1007318

67. Partridge L, Gems D, Withers DJ. Sex and Death: What Is the Connection? Cell. 2005;120(4):461–472. doi:10.1016/j.cell.2005.01.026

68. Catterson JH, Knowles-Barley S, James K, Heck MMS, Harmar AJ, Hartley PS. Dietary Modulation of Drosophila Sleep-Wake Behaviour. PLOS ONE. 2010;5(8):e12062. doi:10.1371/journal.pone.0012062

69. Wang YT, Therriault J, Servaes S, et al. Sex-specific modulation of amyloid-β on tau phosphorylation underlies faster tangle accumulation in females. Brain. 2024;147(4):1497–1510. doi:10.1093/brain/awad397

70. Chen Y, Yu Y. Tau and neuroinflammation in Alzheimer’s disease: interplay mechanisms and clinical translation. Journal of Neuroinflammation. 2023;20(1):165. doi:10.1186/s12974-023-02853-3

71. Yue M, Hanna A, Wilson J, Roder H, Janus C. Sex difference in pathology and memory decline in rTg4510 mouse model of tauopathy. Neurobiology of Aging. 2011;32(4):590–603. doi:10.1016/j.neurobiolaging.2009.04.006

72. Kounatidis I, Chtarbanova S, Cao Y, et al. NF-κB Immunity in the Brain Determines Fly Lifespan in Healthy Aging and Age-Related Neurodegeneration. Cell Reports. 2017;19(4):836–848. doi:10.1016/j.celrep.2017.04.007

73. Shukla AK, Spurrier J, Kuzina I, Giniger E. Hyperactive Innate Immunity Causes Degeneration of Dopamine Neurons upon Altering Activity of Cdk5. Cell Reports. 2019;26(1):131–144.e4. doi:10.1016/j.celrep.2018.12.025

74. Anderson EN, Morera AA, Kour S, et al. Traumatic injury compromises nucleocytoplasmic transport and leads to TDP-43 pathology. Verstreken P, Zoghbi HY, Yamamoto S, eds. eLife. 2021;10:e67587. doi:10.7554/eLife.67587

75. Inequalities in dementia. Dementia Statistics Hub. Accessed May 2, 2026. https://dementiastatistics.org/perceptions-and-inequalities/inequalities/

76. Buckley RF, Mormino EC, Rabin JS, et al. Sex Differences in the Association of Global Amyloid and Regional Tau Deposition Measured by Positron Emission Tomography in Clinically Normal Older Adults. JAMA Neurol. 2019;76(5):542–551. doi:10.1001/jamaneurol.2018.4693

77. Gozes I, Shapira G, Lobyntseva A, Shomron N. Unexpected gender differences in progressive supranuclear palsy reveal efficacy for davunetide in women. Transl Psychiatry. 2023;13(1):319. doi:10.1038/s41398-023-02618-9

78. Bayram E, Carter DJ, Aslam S, Forbes E, Holden SK. Sex differences for clinical presentations and co-pathologies in four-repeat tauopathies. Biol Sex Differ. Published online April 3, 2026. doi:10.1186/s13293-026-00899-5

79. Nebel RA, Aggarwal NT, Barnes LL, et al. Understanding the impact of sex and gender in Alzheimer’s disease: A call to action. Alzheimers Dement. 2018;14(9):1171–1183. doi:10.1016/j.jalz.2018.04.008

80. Klein SL, Flanagan KL. Sex differences in immune responses. Nat Rev Immunol. 2016;16(10):626–638. doi:10.1038/nri.2016.90

81. Altmann A, Tian L, Henderson VW, Greicius MD, Alzheimer’s Disease Neuroimaging Initiative Investigators. Sex modifies the APOE-related risk of developing Alzheimer disease. Ann Neurol. 2014;75(4):563–573. doi:10.1002/ana.24135

82. Blumenfeld J, Yip O, Kim MJ, Huang Y. Cell type-specific roles of APOE4 in Alzheimer disease. Nat Rev Neurosci. 2024;25(2):91–110. doi:10.1038/s41583-023-00776-9

83. Holmes BB, DeVos SL, Kfoury N, et al. Heparan sulfate proteoglycans mediate internalization and propagation of specific proteopathic seeds. Proc Natl Acad Sci U S A. 2013;110(33):E3138–3147. doi:10.1073/pnas.1301440110

84. Blair LJ, Sabbagh JJ, Dickey CA. Targeting Hsp90 and its co-chaperones to treat Alzheimer’s disease. Expert Opin Ther Targets. 2014;18(10):1219–1232. doi:10.1517/14728222.2014.943185

85. Landis GN, Hilsabeck TAU, Bell HS, et al. Mifepristone Increases Life Span of Virgin Female Drosophila on Regular and High-fat Diet Without Reducing Food Intake. Front Genet. 2021;12. doi:10.3389/fgene.2021.751647

86. Li Q, Stavropoulos N. Evaluation of Ligand-Inducible Expression Systems for Conditional Neuronal Manipulations of Sleep in Drosophila. G3 Genes|Genomes|Genetics. 2016;6(10):3351–3359. doi:10.1534/g3.116.034132

87. Aqsa, Sarkar S. Age dependent trans-cellular propagation of human tau aggregates in Drosophila disease models. Brain Research. 2021;1751:147207. doi:10.1016/j.brainres.2020.147207

88. Bankapalli K, Thomas RE, Vincow ES, Milstein G, Fisher LV, Pallanck LJ. A Drosophila model for mechanistic investigation of tau protein spread. Dis Model Mech. 2024;17(9):dmm050858. doi:10.1242/dmm.050858

89. Johnson CE, Duncan MJ, Murphy MP. Sex and Sleep Disruption as Contributing Factors in Alzheimer’s Disease. Journal of Alzheimer’s Disease. 2024;97(1):31–74. doi:10.3233/JAD-230527

